# Using the residual bootstrap to quantify uncertainty in mean apparent propagator MRI

**DOI:** 10.1101/295667

**Authors:** Xuan Gu, Anders Eklund, Evren Özarslan, Hans Knutsson

## Abstract

Estimation of noise-induced variability in MAP-MRI is needed to properly characterize the amount of uncertainty in quantities derived from the estimated MAP-MRI coefficients. Bootstrap metrics, such as the standard deviation, provides additional valuable diffusion information in addition to common MAP-MRI parameters, and can be incorporated in MAP-MRI studies to provide more extensive insight. To the best of our knowledge, this is the first paper to study the uncertainty of MAP-MRI derived metrics. The noise variability of quantities of MAP-MRI have been quantified using the residual bootstrap, in which the residuals are resampled using two sampling schemes. The residual bootstrap method can provide empirical distributions for MAP-MRI derived quantities, even when the exact distributions are not easily derived. The residual bootstrap methods are applied to SPARC data and HCP-MGH data, and empirical distributions are obtained for the zero-displacement probabilities. Here, we compare and contrast the residual bootstrap schemes using all shells and within the same shell. We show that residual resampling within each shell generates larger uncertainty than using all shells for the HCP-MGH data. Standard deviation and quartile coefficient maps of the estimated variability are provided.

## 1 Introduction

Mean apparent propagator (MAP) MRI is a diffusion-weighted MRI framework for accurately characterizing and quantifying anisotropic diffusion properties at large as well as small levels of diffusion sensitivity (Özarslan et al., 2013). Consequently, it has been demonstrated that MAP-MRI can capture intrinsic nervous tissue features (Avram et al., 2016; Fick et al., 2016; Özarslan et al., 2013). Some novel features of the diffusion can be characterized by MAP-MRI, including the return-to-origin probability (RTOP), return-to-plane probability (RTPP), and the return-to-axis probability (RTAP). With the assumptions that the gradients are infinitely short and the diffusion time is sufficiently long, these MAP-MRI derived metrics describe the mean pore volume and cross-sectional area for a population of isolated pores. It is, however, not clear how high the uncertainty is for these measures.

Bootstrap is a non-parametric statistical technique, based on data resampling, used to quantify uncertainties of parameters (Efron, 1992). Bootstrap has been widely used in diffusion tensor imaging (DTI) to study uncertainty associated with DTI parameter estimation (Chung et al., 2006; Heim et al., 2004; Pajevic and Basser, 2003; Vorburger et al., 2016, 2012; Yuan et al., 2008). The repetition bootstrap method requires multiple measurements per gradient direction to perform resampling (Heim et al., 2004). For most clinical and research applications, it is more interesting to obtain more gradient directions, since diffusion parameter estimation can be more robust with high angular resolution diffusion imaging (HARDI). To be able to use bootstrap for diffusion data with multi gradient directions, instead of repetitions of the same direction, residual bootstrap can be used with the assumption that the error terms have constant variance. The wild bootstrap (Wu, 1986) is suited when the data are heteroscedastic (i.e., non-constant variance). The sensitivity of the MAP-MRI metrics on noise, experimental design, and the estimation of the scaling tensor has been investigated in (Hutchinson et al., 2017).

Implementation of the repetition bootstrap, residual bootstrap and wild bootstrap have already been reported for DTI (Chung et al., 2006; Efron, 1992; Heim et al., 2004; Vorburger et al., 2016, 2012; Yuan et al., 2008) while there are no reports of bootstrap for MAP-MRI. A possible explanation is the higher computational complexity of MAP-MRI. The explanation is that DTI has been used for a long time, while MAP-MRI is a rather new framework. We use a model-based resampling technique, the residual bootstrap, that may be applied to the residuals from the linear regression model used to fit MAP-MRI in each voxel. The residual bootstrap is specifically designed to work when data are homoscedastic; that is, when the variance of the errors is constant for all observations. For the standard diffusion tensor model the non-constant variance comes from the log-transform of the diffusion signal, but no log-transform is applied in MAP-MRI.

This paper investigates the ability of model-based resampling, in particular, the residual bootstrap, to provide reasonable estimates of variability for MAP-MRI derived quantities for physical phantom data (SPARC) (Ning et al., 2015) and for human brain data (HCP-MGH) (Van Essen et al., 2013). Two sampling schemes, residual sampling using all shells and within each shell, are used to produce uncertainty maps. Comparisons are made between the two sampling schemes, between data with different b-values, and also for different healthy subjects. To the best of our knowledge, this is the first paper to study the uncertainty of MAP-MRI derived metrics.

## 2 Theory

### 2.1 MAP-MRI

The three-dimensional q-space diffusion signal attenuation *E*(**q**) is expressed in MAP-MRI as

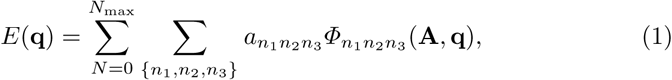

where *Φ*_*n*_1_ *n*_2_ *n*_2__ (**A**, **q**) are related to Hermite basis functions and depend on the second-order tensor **A** and the **q**-space vector **q**. The non-negative indices *n*^*i*^ are the order of Hermite basis functions which satisfy the condition *n*_1_ + *n*_2_ + *n*_3_ = *N*. The q-space vector **q** is defined as **q** = *γδ***G**/2*π*, where *γ* is the gyromagnetic ratio, *δ* is the diffusion gradient duration, and **G** determines the gradient strength and direction. The propagator is the three-dimensional inverse Fourier transform of *E*(**q**), and can be expressed as

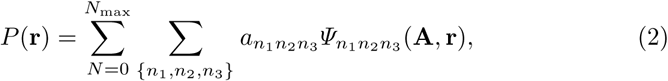

where *Ψ*_*n*_1_ *n*_2_ *n*_2__ (**A**, **r**) are the corresponding basis functions in displacement space **r**. The number of coefficients for MAP-MRI is given by

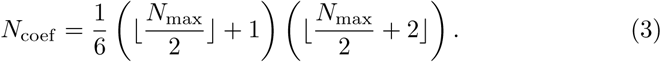

**A** can be taken to be the covariance matrix of displacement, defined as

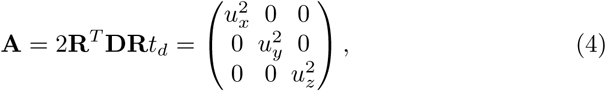

where **R** is the transformation matrix whose columns are the eigenvectors of the standard diffusion tensor **D**, and *t*_d_ is the diffusion time. The MAP-MRI basis functions, *Φ*_*n*_1_ *n*_2_ *n*_2__ (**A**, **q**) in **q**-space and *Ψ*_*n*_1_ *n*_2_ *n*_2__ (**A**, **r**) in displacement **r**-space, are given by

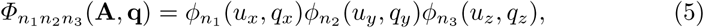

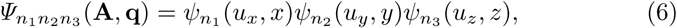

with (Özarslan et al., 2008)

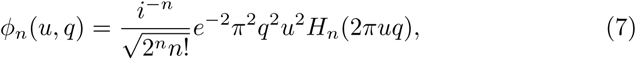

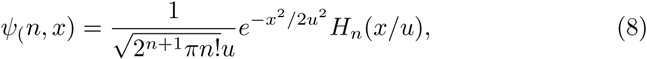

where *H*_*n*_(*x*) is the *n*th order Hermite polynomial. Equation 1 can be written in matrix form as

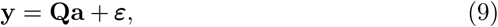

where **y** is a vector of *T* signal values, **Q** is a *T* × *N*_coef_ design matrix formed by the basis functions *Φ*_*n*_1_ *n*_2_ *n*_2__ (**A**, **q**), **a** contains the parameters to estimate, and ***ε*** is the error. The coefficients **a** can be obtained by solving the following quadratic minimization problem,

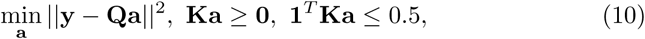

where **0** and **1** are vectors with elements 0 and 1, respectively. The rows of the constraint matrix **K** are the basis functions *Ψ*_*n*_1_ *n*_2_ *n*_2__ (**A**, **q**) evaluated on a uniform Cartesian grid in the positive *z* half space. The first constraint enforces non-negativity of the propagator, and the second one limits the integral of the probability density to a value no greater than 1.

Zero displacement probabilities include the return-to-origin probability (RTOP), and its variants in 1D and 2D: the return-to-plane probability (RTPP), and the return-to-axis probability (RTAP), respectively. Return-to-origin-probability, *P*(**r**), is the probability for water molecules to undergo no net displacement. In terms of MAP-MRI coefficients through the expression it is defined as

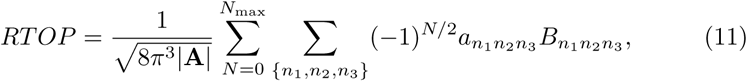

where

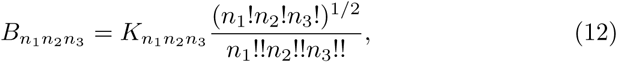

and *K*_*n*_1_ *n*_2_ *n*_2__ = 1 if *n*_1_, *n*_2_ and *n*_3_ are all even and 0 otherwise. If we consider a population of isolated pores, with the assumptions that the diffusion gradients are infinitesimally short and the diffusion time is sufficiently long, it can be shown that (Özarslan et al., 2013)

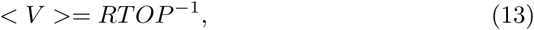

which indicates that the reciprocal of the RTOP is the statistical mean pore volume.

RTAP indicates the probability density for molecules to return to the axis determined by the principal eigenvector of the A-matrix. On the other hand, RTPP represents the likelihood for no net displacement along this axis. If the principal eigenvector is along the *x*-axis, RTAP and RTPP are given by

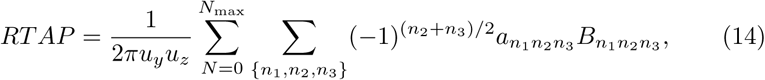

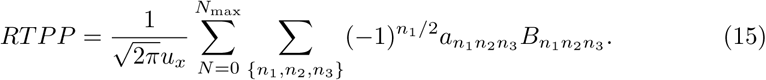

### 2.2 Bootstrap

Repetition (regular) bootstrap requires multiple measurements per gradient direction, and for each gradient direction the measurements are sampled with replacement over-and-over again to characterize the uncertainty of the diffusion derived metrics (Heim et al., 2004). However, nowadays it becomes clinically more feasible to have scan protocols with a large number of gradient directions, instead of having more than one measurement per direction.

Alternatives to repetition bootstrap are model-based bootstrap approaches, such as the residual bootstrap and the wild bootstrap. Residual bootstrap relies on the assumption that the residuals are independent and identically distributed (e.g. constant variance); the sample diffusion data are generated by randomly sampling with replacement from the residuals. Wild bootstrap is designed for heteroscedastic data, that is when the variance of the errors is not constant. In the case of the diffusion tensor model(Basser et al., 1994), it is known that the log-transform leads to non-constant variance (Wegmann et al., 2017). Therefore, the residuals are weighted by the heteroscedasticity consistent covariance matrix estimator and random samples are drawn from the auxiliary distribution (Davidson and Flachaire, 2008).

Under the assumption that *ε* is a vector of independent and identically distributed (IID) values with zero mean, the linear regression would be termed homoscedastic and we would be able to randomly sample with replacement from the residual, taking the form

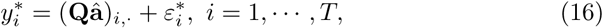

where (**Qâ**)_*i*_,. is the *i*th row of the product of **Q** and **â**, **â** is the solution of the quadratic minimization problem in Equation 10, and 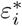 is a random sample from the residuals of the original regression model 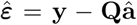. The mean squared error (MSE) can be used to quantify the quality of the signal reconstruction, according to

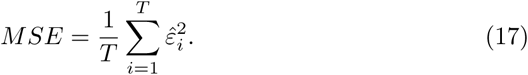

A common test for heteroscedasticity is the White test (White, 1980). A White test can be performed using a second regression, according to

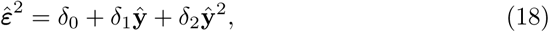

where ŷ is the predicted diffusion signal. It is known that a higher b-value will result in diffusion data with a lower signal-to-noise-ratio (SNR). It has been reported that the SNR issue becomes important at higher b-values, when fitting the data to a diffusion model (Gu et al., 2017). Hence it is more reasonable to resample the residuals from the same b-value group (diffusion shell). We will refer the bootstrap scheme as Scheme A when sampling using all shells, and Scheme B when sampling within each shell separately.

Solving the quadratic minimization problem for 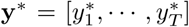 will produce a bootstrap estimate of the coefficients **a**\*. Repeating these steps for some fixed large number *N*_*B*_, resampling and estimation, builds up a collection of co-efficients 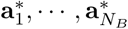 called the bootstrap distribution, from which some MAP-MRI scalar indices can be calculated. Summary statistics from this empirical distribution can be used to describe the original parameter estimate. Here the sample statistic 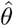 is an estimate of the true unknown *θ* (such as the noise-free RTOP of the voxel) using the original data **y**, and 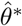 is the bootstrap replication of 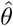. The bootstrap-estimated standard error of 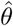 is simply the standard deviation of the *N*_*B*_ replications, i.e.,

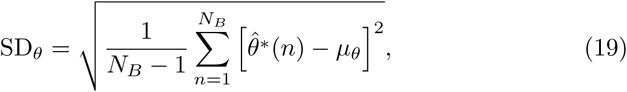

where 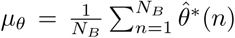. The Quartile coefficient (QC) is a dimensionless measure of dispersion which can be computed using the first (*Q*1) and third (*Q*3) quartiles (Bonett, 2006), according to

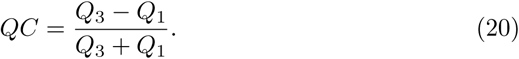

In this paper we use QC for comparing the dispersion of parameters.

## 3 Data and Methods

### 3.1 SPARC phantom data

We use data from the Sparse Reconstruction Challenge for Diffusion MRI (SPARC dMRI) hosted at the 2014 CDMRI workshop on computational diffusion MRI (Ning et al., 2015). The data were acquired from a physical phantom with known fiber configuration. The phantom is made of polyfil fibers of 15 *µ*m diameter (Moussavi-Biugui et al., 2011). It provides a mask to indicate the number of fiber bundles crossing in each voxel. In two-fiber voxels, the fiber bundles are crossing at a 45 degree angle with isotropic diffusion outside. The voxels that are masked by 0 have no fibers and are not considered. Three sets of data are acquired with b-values of 1000, 2000, and 3000 s/mm^2^, using 20, 30 and 60 gradient directions per shell for the three datasets respectively (hereinafter referred to as SPARC-20, SPARC-30 and SPARC-60). The gold-standard data was obtained by acquiring 81 gradient directions at b-values of 1000, 2000, 3000, 4000 and 5000 s/mm^2^ averaged over 10 repetitions, resulting in 405 measurements (hereinafter referred to as SPARC-Gold). All datasets include one measurement with *b*_0_. The data has dimension 13 × 16 × 406 and resolution 2 × 2 × 7 mm. The diffusion time and pulse separation time are *δ* = *∆* = 62ms.

### 3.2 Human Connectome Project MGH adult diffusion data

We use the MGH adult diffusion dataset from the Human Connectome Project (HCP) (Van Essen et al., 2013). Data were collected from 35 healthy adults scanned on a customized Siemens 3T Connectom scanner with 4 different b-values (1000, 3000, 5000 and 10,000 s/mm^2^). The data has already been preprocessed for gradient nonlinearity correction, motion correction and eddy current correction (Glasser et al., 2013). The data consists of 40 non-diffusion weighted volumes (b = 0), 64 volumes for b = 1000 and 3000 s/mm^2^, 128 volumes for b = 5000 s/mm^2^ and 256 volumes for b = 10, 000 s/mm^2^, which yields 552 volumes of 140 × 140 × 96 voxels with an 1.5 mm isotropic voxel size. The diffusion time and pulse separation time are *δ* = 12.9 ms and *∆* = 21.8 ms. The HCP-MGH data also contains high-resolution T1 images of 256 × 256 × 276 voxels with an 1.0 mm isotropic voxel size.

Data used in the preparation of this work were obtained from the Human Connectome Project (HCP) database (https://ida.loni.usc.edu/login.jsp). The HCP project (Principal Investigators: Bruce Rosen, M.D., Ph.D., Martinos Center at Massachusetts General Hospital; Arthur W. Toga, Ph.D., University of Southern California, Van J. Weeden, MD, Martinos Center at Massachusetts General Hospital) is supported by the National Institute of Dental and Craniofacial Research (NIDCR), the National Institute of Mental Health (NIMH) and the National Institute of Neurological Disorders and Stroke (NINDS). HCP is the result of efforts of co-investigators from the University of Southern California, Martinos Center for Biomedical Imaging at Massachusetts General Hospital (MGH), Washington University, and the University of Minnesota.

### 3.3 Methods

Diffusion tensor fitting, MAP-MRI fitting and bootstrap sampling were implemented using C++. The initial tensor fitting was performed with data with b-values less than 2000 s/mm^2^ using weighted least squares. To impose the constraint of positivity of the propagator, we sample *P* (**r**) in a 21 × 21 × 11 grid, resulting in 4851 points. Here the last dimension is only sampled on its positive axis as the propagator is antipodally symmetric. We use the Gurobi Optimizer (Gurobi Optimization, 2016) to solve the quadratic optimization problem. The Open Multi-Processing (OpenMP) (Dagum and Menon, 1998) framework is used to run the analysis for many voxels in parallel. MAP-MRI fitting and bootstrap sampling are computationally expensive, due to the large number of MAP coefficients, constraints in the quadratic minimization problem and repeating the analysis 500-5000 times. We use a computer with 512 GB RAM and two Intel(R) Xeon(R) E5-2697 2.30 GHz CPUs. Each of the two CPUs has 18 cores (36 threads), which makes it possible to run the analysis for 72 voxels in parallel.

In order to perform a voxel-level group comparisons of diffusion-derived metric maps, the diffusion data must be transformed to a standard space. The transformation between MNI standard space and diffusion space is achieved in three separate steps. First, the non-diffusion volume is registered to the T1 volume using the *FSL* (Jenkinson et al., 2012) function *epi_reg*. Second, the T1 volume is non-linearly registered to the MNI152 T1 2mm template using the *FSL* function *fnirt* (Andersson et al., 2007). Third, the two transformations were combined, to transform the diffusion data to MNI space. The statistics analysis is performed in MATLAB (R2016b, The MathWorks, Inc., Natick, Massachusetts, United States).

## 4 Results

### 4.1 SPARC

Results of the SPARC-Gold data will be used as a comparison reference for the following study on SPARC-20, SPARC-30 and SPARC-60. Figure 1 shows scalar maps of the fiber bundles mask, fractional anisotropy (FA), mean diffusivity (MD) and RTOP^1/3^. The values in the fiber bundles mask indicate the number of fiber bundles in each voxel. The voxels masked by 0 are referred as empty area. The construction of the physical phantom is described in (Moussavi-Biugui et al., 2011). The MD clearly shows different diffusivities in two-fiber areas (4.78 *±* 0.63 × 10^−4^ mm^2^/s) and single-fiber areas (7.51 *±* 0.67 × 10^−4^ mm^2^/s). In single-fiber and crossing-fiber areas the FA shows very similar values (0.82 *±* 0.03 and 0.85 *±* 0.01, respectively). Finally, for the *N*_max_ of 6 for MAP-MRI, we find fairly consistent RTOP^1/3^ values in both single-fiber (81.0±7.87 m^−1^) and two-fiber (111.7 *±* 11.1 m^−1^) areas and higher in the two-fiber areas. Also, it can be noticed that RTOP^1/3^ in two-fiber area shows a larger degree of variation (larger standard deviation).

**Fig. 1.**
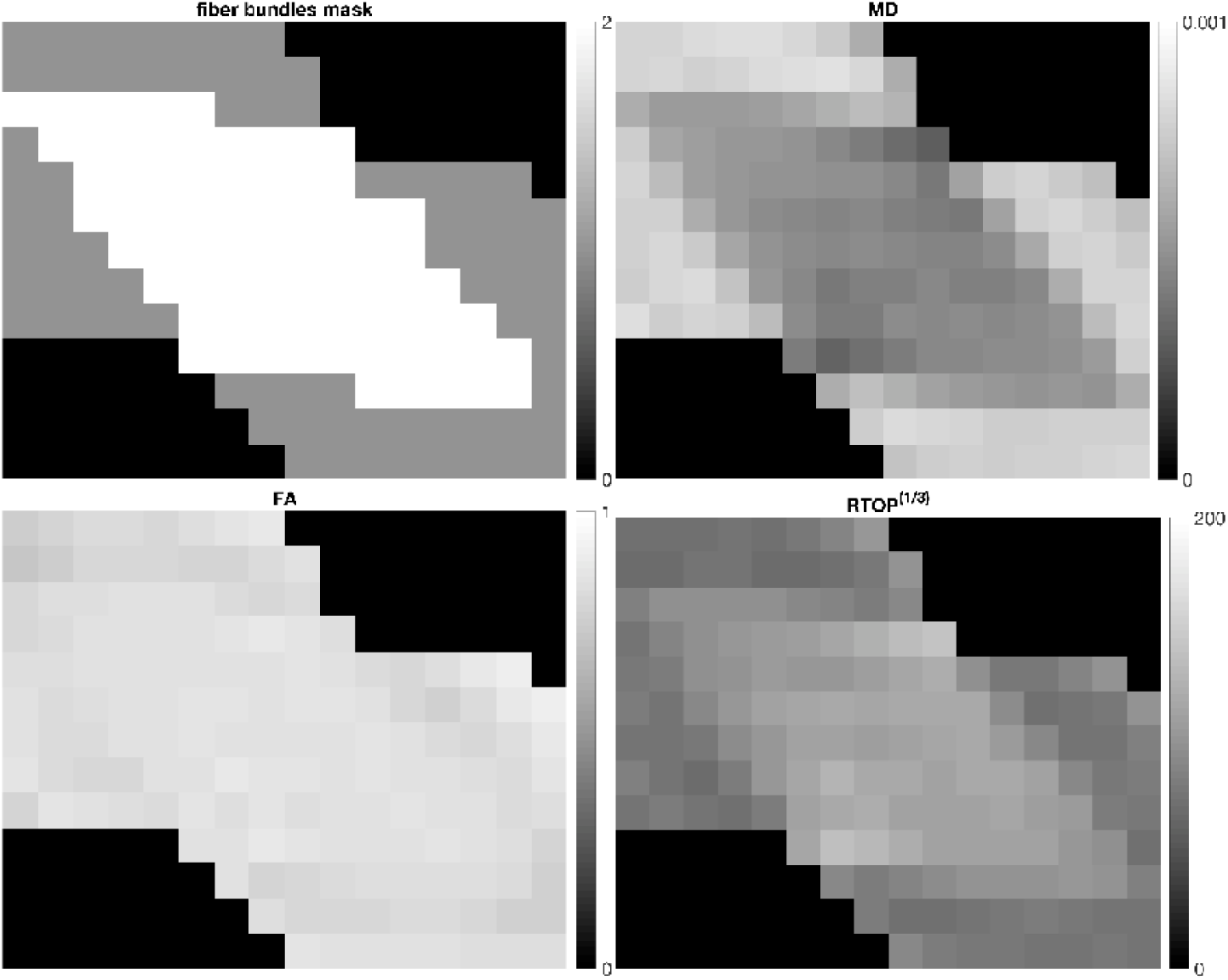
Top left: the fiber bundles mask, top right: Mean Diffusivity (MD) (mm^2^/s), bottom left: Fractional Anisotropy (FA), bottom right: RTOP^1/3^ (mm^−1^) for SPARC-Gold. It can be seen that MD differs for different fiber configurations, while FA is more or less constant in fibrous areas. The RTOP^1/3^ shows lower values in single-fiber areas, but higher in two-fiber voxels.

To study the effect of the MAP-MRI *N*_max_ on signal reconstruction, we compare the signal reconstruction quality over different *N*_max_s for SPARC-Gold. Figure 2 shows the original diffusion signal, the MAP-MRI fitted signal and the residuals for an unidirectional voxel (1,1) and a bidirectional voxel (8,8) from SPARC-Gold. For comparison, we normalized the signal with the first volume (*b*_0_). For higher *N*_max_s both voxels have a lower error. There is a substantially lower error in single-fiber voxels than two-fiber voxels. Larger *N*_max_s (more basis functions) provide a better characterization of the diffusion signal. The amount of additional detail in the reconstruction of the propagator does not increase beyond a certain point. The trade-off between the level of detail in the propagator estimation, the amount of data acquired and the computation time is important. It is reported that including terms up to order 6 was found to yield a sufficient level of detail in propagators from diverse brain regions (Fick et al., 2016). In this paper all further analysis of MAP-MRI parameters described in this paper use *N*_max_ = 8, if not specified otherwise.

**Fig. 2.**
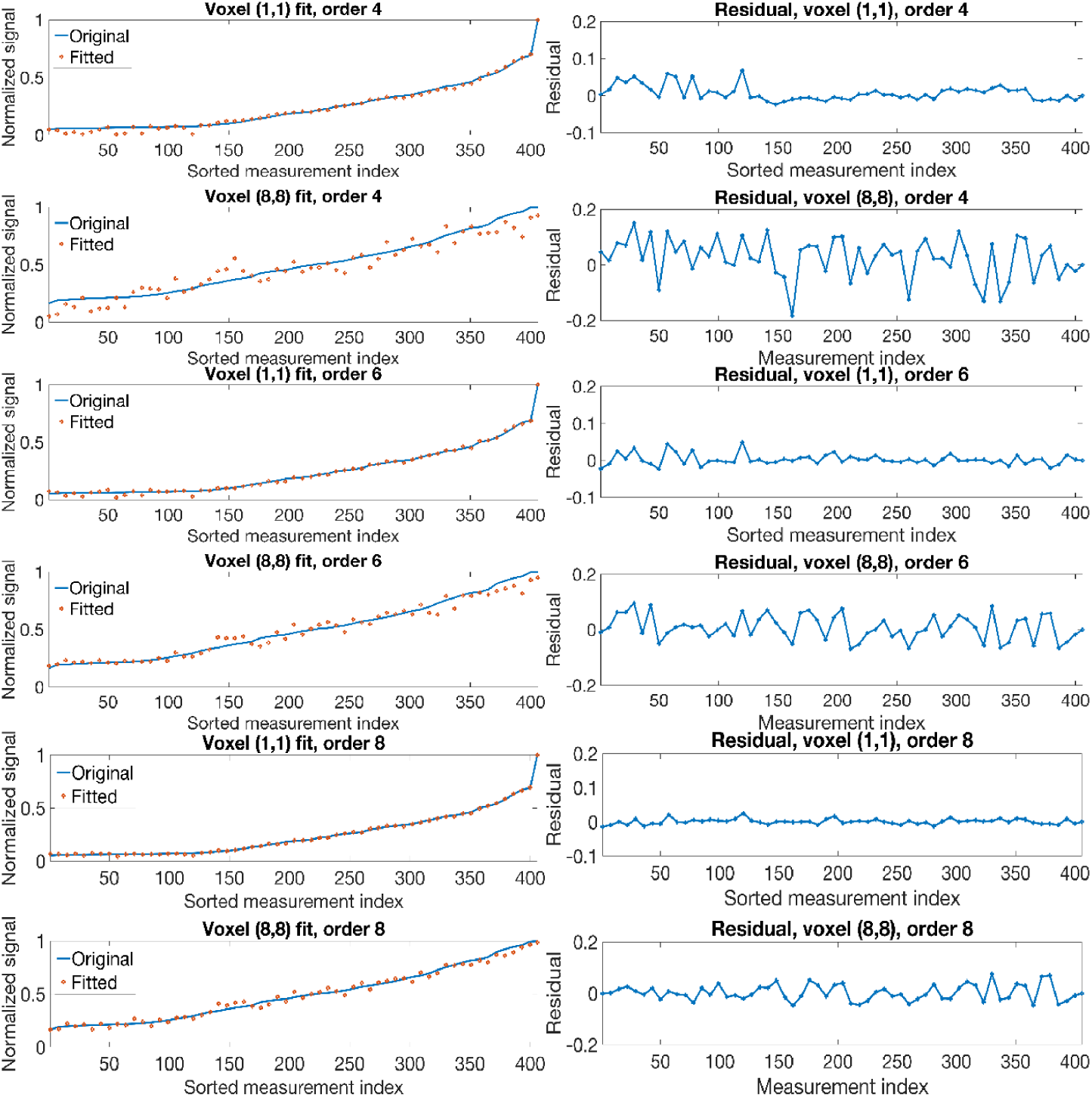
Normalized diffusion signals, MAP-MRI fitted signals and residuals for an unidirectional voxel (1,1) and a bidirectional voxel (8,8) from SPARC-Gold, using *N*_max_ 4, 6 and 8. Measurement indices are sorted in an ascending order of the signal intensities. The 406 samples represent diffusion data from *b* = 0, 1000, 2000, 3000, 4000, 5000 mm^2^/s

To study the quality of the MAP-MRI signal fitting, we focus on two voxels; one unidirectional voxel (1,1) and one bidirectional voxel (8,8). The mean squared error (MSE) and the computation time (using 30 CPU threads) of the signal recovery for the two voxels is summarized in Figure 3. Please note that the computation time plotted is for the SPARC-30 data, in order to compare with the results in (Fick et al., 2016). It can be noted that the quality of signal fitting for the single-fiber voxel (1,1) quickly reaches the optimum when the *N*_max_ is larger than 4. For the two-fiber voxel (8,8), no substantial differences in the signal fitting can be observed when the *N*_max_ is beyond 6. The number of MAP-MRI coefficients to be estimated in each voxel for the *N*_max_ of 2 to 10 is 7, 22, 50, 95, 161, respectively. With the help of OpenMP and the Gurobi Optimizer, we are able to perform the MAP-MRI fitting for SPARC-30 using a *N*_max_ of 10 within 15 seconds, which is 17 times faster than its counterpart in (Fick et al., 2016).

**Fig. 3.**
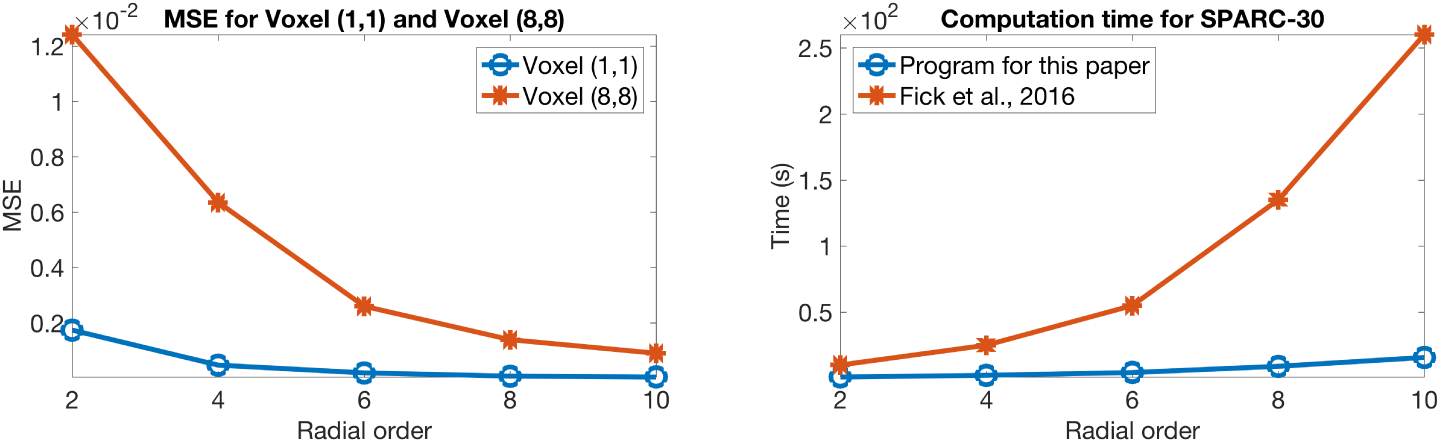
Left: the mean squared error of the reconstructed signal with respect to the diffusion signal for an unidirectional voxel(1,1) and a bidirectional voxel(8,8) from SPARC-Gold. Right: the computation time in seconds for two implementations of MAP-MRI.

Figure 4 show the standard deviation of RTOP^1/3^ for SPARC-20, SPARC-30, SPARC-60 and SPARC-Gold, using the two bootstrap sampling schemes and 500 bootstrap samples. The standard deviation values are directly proportional to the RTOP values. For all datasets the standard deviation maps exhibit minor differences for the two sampling schemes. A reduction in the standard deviation is observed with an increasing number of diffusion measurements and gradient directions from SPARC20 to SPARC-Gold. This is explained by the fact that the data acquisition protocols with a larger number of diffusion measurements and gradient directions will improve the signal fitting quality, which reduces the variability of the estimated RTOP values. As shown in Figure 5, it is notable that the mean standard deviation of the two-fiber area demonstrates a downward trend from SPARC-20 to SPARC-60, as the number of measurements increased from 41 to 61 and 121, using the same gradient strength. With 406 measurements and b-values of 1000 to 5000, the mean standard deviation of the two-fiber area for SPARC-Gold is dramatically decreased. The single-fiber area is not affected by the number of measurements.

**Fig. 4.**
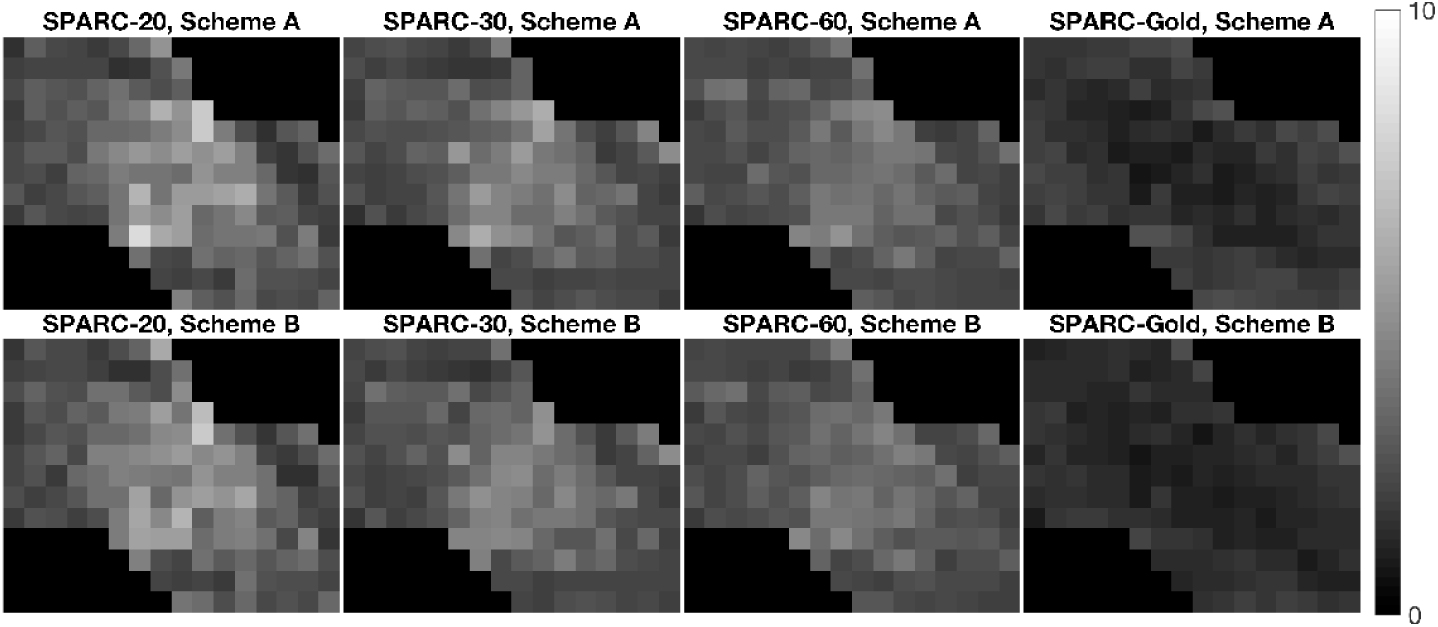
Standard deviation of the RTOP^1/3^ for SPARC-20, SPARC-30, SPARC-60 and SPARC-Gold, using two bootstrap sampling schemes and 500 bootstrap samples. The standard deviation is clearly lower for SPARC-Gold. Scheme A: sampling using all shells, Scheme B: sampling within each shell.

**Fig. 5.**
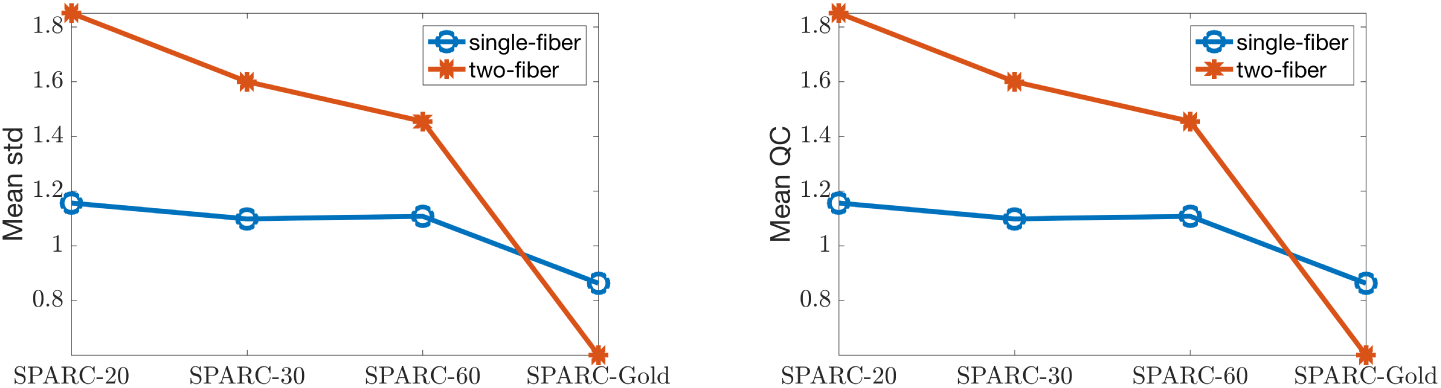
Mean standard deviation (left) and mean QC (right) of the RTOP^1/3^ for single-fiber voxels and two-fiber voxels in SPARC-20, SPARC-30, SPARC-60 and SPARC-Gold, using bootstrap Scheme A (sampling using all shells) and 500 samples.

Figure 6 shows the QC of the RTOP^1/3^ for SPARC-20, SPARC-30, SPARC-60 and SPARC-Gold, using two bootstrap sampling schemes and 500 samples. The mean quartile coefficients for SPARC-Gold show less variability, and this is to be expected since SPARC-Gold is based on more gradient directions and more measurements. We can see that one potential source of variability comes from the bootstrap sampling methodology. Data with a larger number of gradient directions produces more stable estimates of MAP-MRI coefficients, from which RTOP is based. Scheme A and B for SPARC-Gold data produce similar estimates of uncertainty. The QC maps from Scheme B show more smooth and consistent results for SPARC-20, SPARC-30 and SPARC-60. It is interesting to note that the QC maps from all sampling schemes for SPARC-Gold are exhibiting larger variability for single-fiber areas, compared to two-fiber areas.

**Fig. 6.**
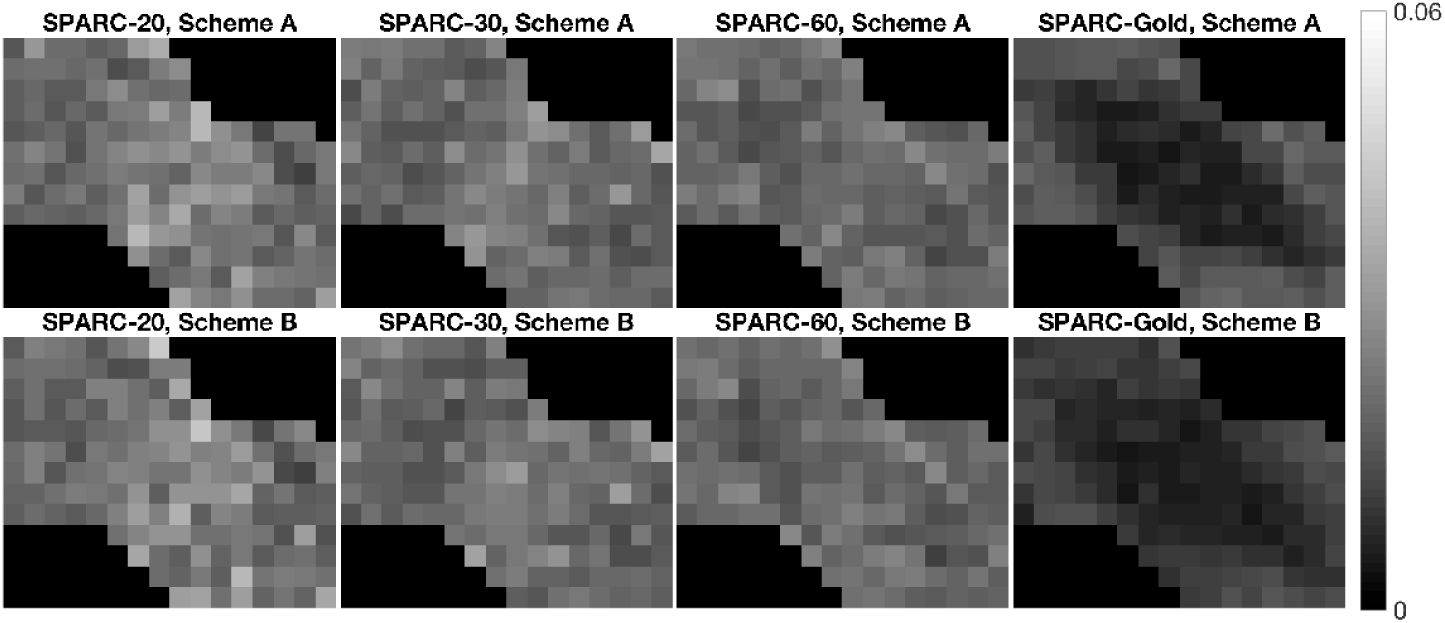
QC of the RTOP^1/3^ for SPARC-20, SPARC-30, SPARC-60 and SPARC-Gold, using two bootstrap sampling schemes and 500 bootstrap samples. Scheme A: sampling using all shells, Scheme B: sampling within each shell.

Figure 7 shows the histograms of RTOP^1/3^ for SPARC-60 using one bootstrap sample and 500 samples. The mean values for single-fiber and two-fiber areas are plotted as vertical lines. With 500 samples, the histogram is able to reflect the distribution in which two patterns can be detected, corresponding to the two fiber configurations.

**Fig. 7.**
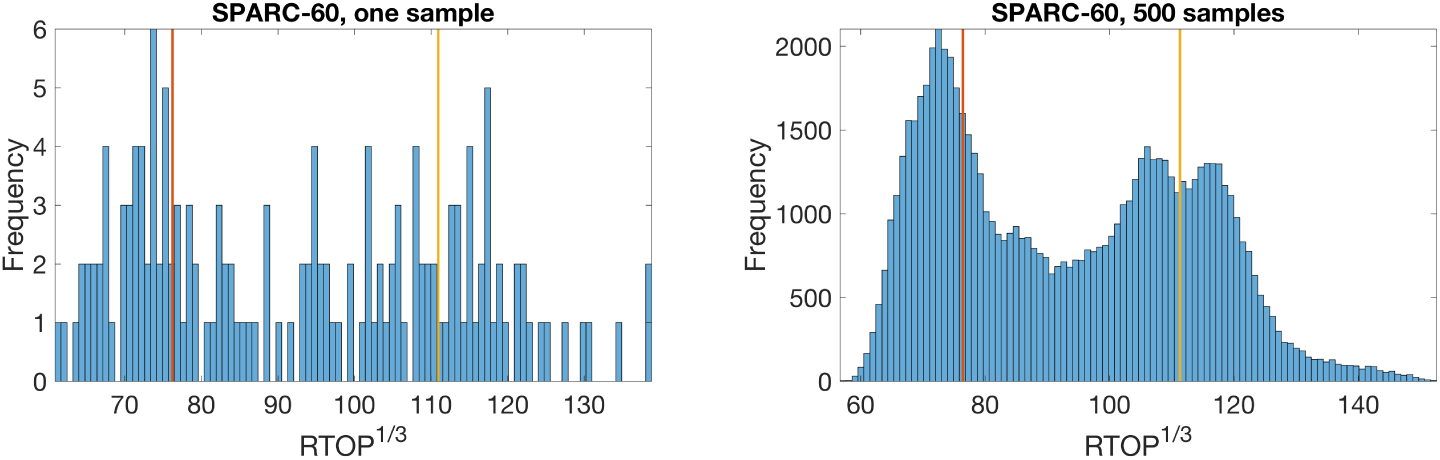
Histograms of the RTOP^1/3^ for all voxels in SPARC-60, using bootstrap Scheme A (sampling using all shells) and 500 bootstrap samples.

Figure 8 and 9 show the histograms of the RTOP^1/3^ of an unidirectional voxel (1,1) and a bidirectional voxel (8,8) for SPARC-20, SPARC-30, SPARC-60 and SPARC-Gold, using two bootstrap sampling schemes and 500 bootstrap samples. It is interesting to note that the means of the RTOP^1/3^ of the unidirectional voxel (1,1) are different for Scheme A and Scheme B.

**Fig. 8.**
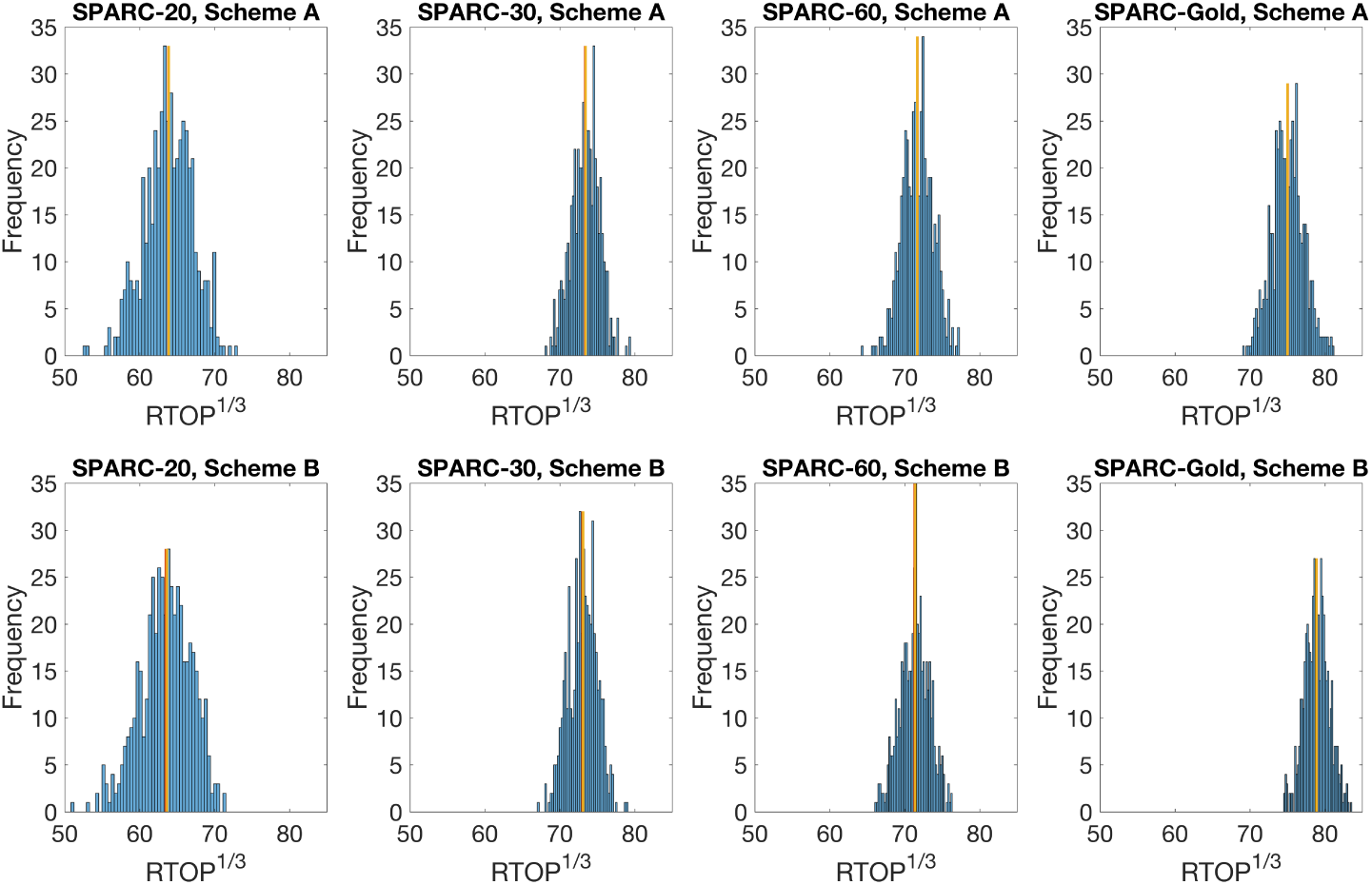
Histograms of the RTOP^1/3^ of an unidirectional voxel (1,1) for SPARC-20, SPARC-30, SPARC-60 and SPARC-Gold, using two bootstrap sampling schemes and 500 bootstrap samples. Scheme A: sampling using all shells, Scheme B: sampling using the same shell.

**Fig. 9.**
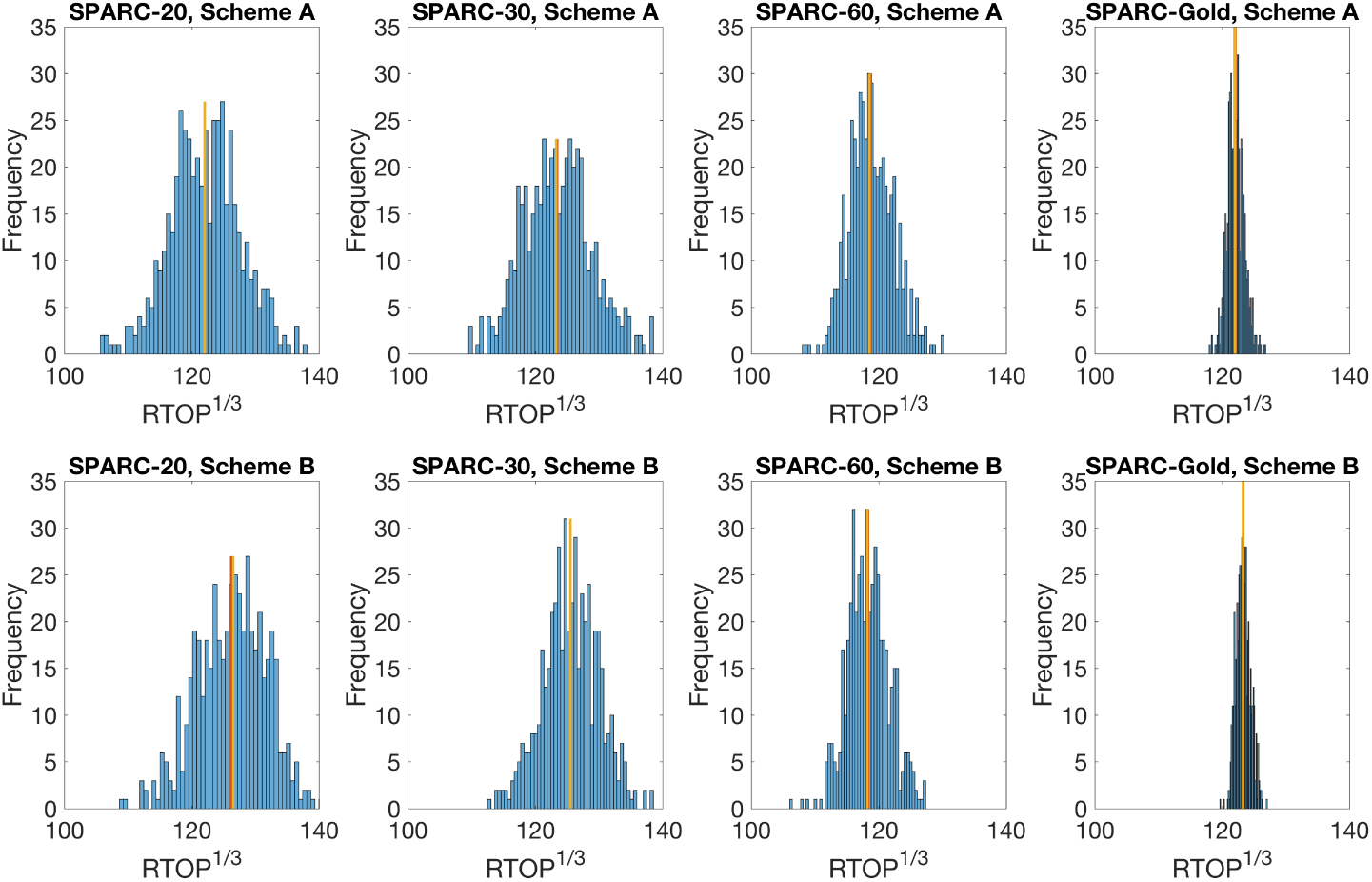
Histograms of the RTOP^1/3^ of a bidirectional voxel (8,8) for SPARC-20, SPARC-30, SPARC-60 and SPARC-Gold, using two bootstrap sampling schemes and 500 bootstrap samples. Scheme A: sampling using all shells, Scheme B: sampling within each shell.

Figure 10 shows the PDF of the RTOP^1/3^ for SPARC-20, SPARC-30, SPARC-60 and SPARC-Gold, using two bootstrap sampling schemes and 500 bootstrap samples. The PDFs of RTOP^1/3^ for SPARC-20, SPARC-30 and SPARC-60 show only minor differences between the two bootstrap sampling schemes since these three datasets were acquired using the same b-values. Both bootstrap sampling schemes show the ability to detect some patterns in the PDF of the RTOP^1/3^ corresponding to the two fiber configurations in the phantom, although with a few notable differences. The SPARC-Gold data does exhibit a different distribution when compared with the other three. It can be inferred that the bootstrap-generated distributions might be related to the b-value settings used in the scans but less affected by the number of measurements.

**Fig. 10.**
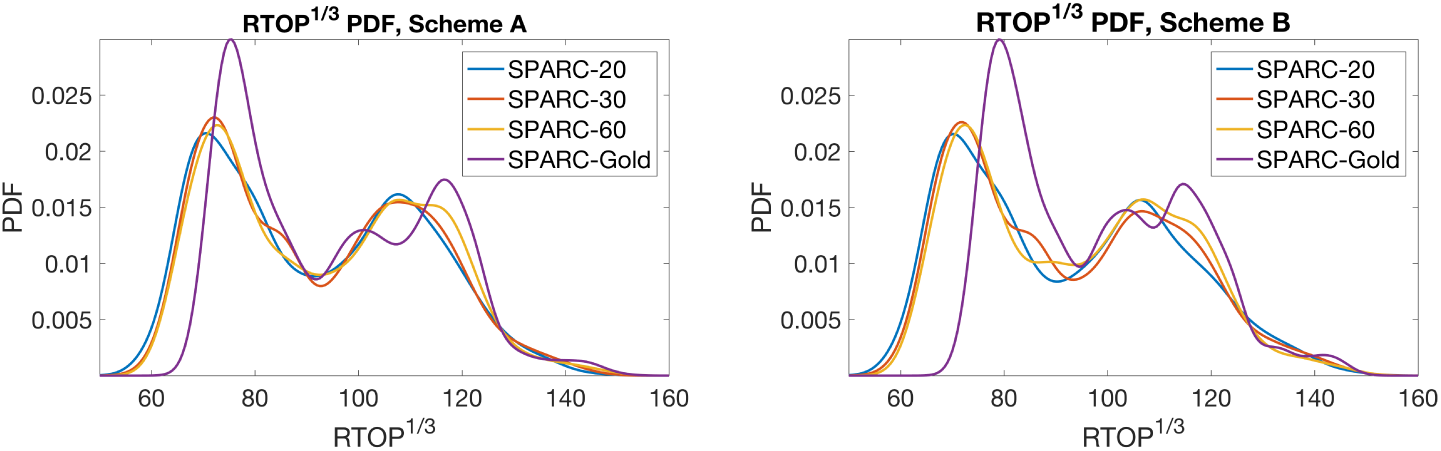
PDF of the RTOP^1/3^ for SPARC-Gold, SPARC-20, SPARC-30 and SPARC-60, using two bootstrap sampling schemes and 500 bootstrap samples. Scheme A: sampling using all shells, Scheme B: sampling within each shell.

Figure 11 shows the boxplots of the RTOP^1/3^ for an unidirectional voxel (1,1) and a bidirectional voxel (8,8) for SPARC-20, SPARC-30, SPARC-60 and SPARC-Gold, using two bootstrap sampling schemes and 500 bootstrap samples. On each box, the central mark indicates the median, and the bottom and top edges of the box indicate the 25th and 75th percentiles *Q*_1_ and *Q*_3_, respectively. The whiskers represent the ranges for the bottom and the top boundaries of the data values, excluding outliers. The boundaries are calculated as *Q*_3_ − 1.5 × (*Q*_3_–*Q*_1_) and *Q*_3_ + 1.5 × (*Q*_3_–*Q*_1_) (McGill et al., 1978), which corresponds to approximately 99.3 percent coverage if the data are normally distributed. The outliers are plotted individually using the ‘+’ symbol beyond the boundaries. Non-skewed data, with a median in the center, indicates that data may be normally distributed. For both voxels, the RTOP distributions for SPARC-20 exhibits a larger amount of variability. For a bidirectional voxel (8, 8), the bootstrap samples of RTOP show much narrower distributions. The boxplots of the RTOP^1/3^ indicate little difference in the two sampling schemes.

**Fig. 11.**
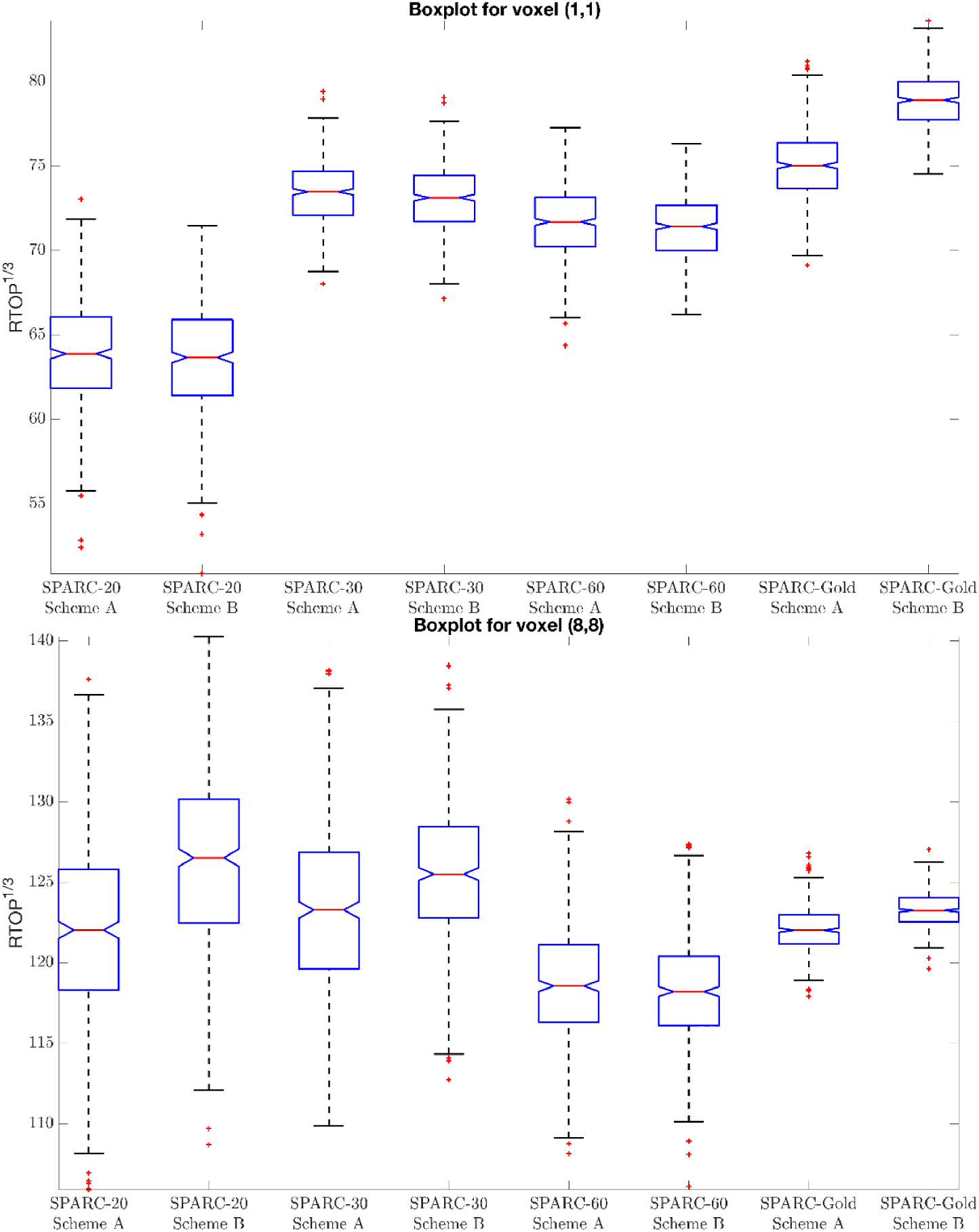
Boxplot of the RTOP^1/3^ for an unidirectional voxel (1,1) (top) and a bidirectional voxel (8,8) (bottom) for SPARC-20, SPARC-30, SPARC-60 and SPARC-Gold, using two bootstrap sampling schemes and 500 bootstrap samples. Scheme A: sampling using all shells, Scheme B: sampling within each shell. The uncertainty is clearly lower for SPARC-Gold

### 4.2 HCP-MGH

In the following section, we present results from HCP-MGH diffusion data.

#### White test

Figure 12 shows examples of MD, FA and RTOP of a slice from subject MGH-1010. Figure 13 shows the results from the White test for the residuals from all shells, and within each shell. The first column demonstrates that most voxels produce a p-value below 0.05 (uncorrected for multiple comparisons) which rejects the null hypothesis of homoskedasticity for the residuals from all shells. Columns 2 to 5 show that 5.1%, 4.9%, 5.3% and 5.0% of the voxels in the brain demonstrate heteroskedasticity, i.e. close to the expected 5% if the null hypothesis is true. These tests show that sampling within each shell should be used.

**Fig. 12.**
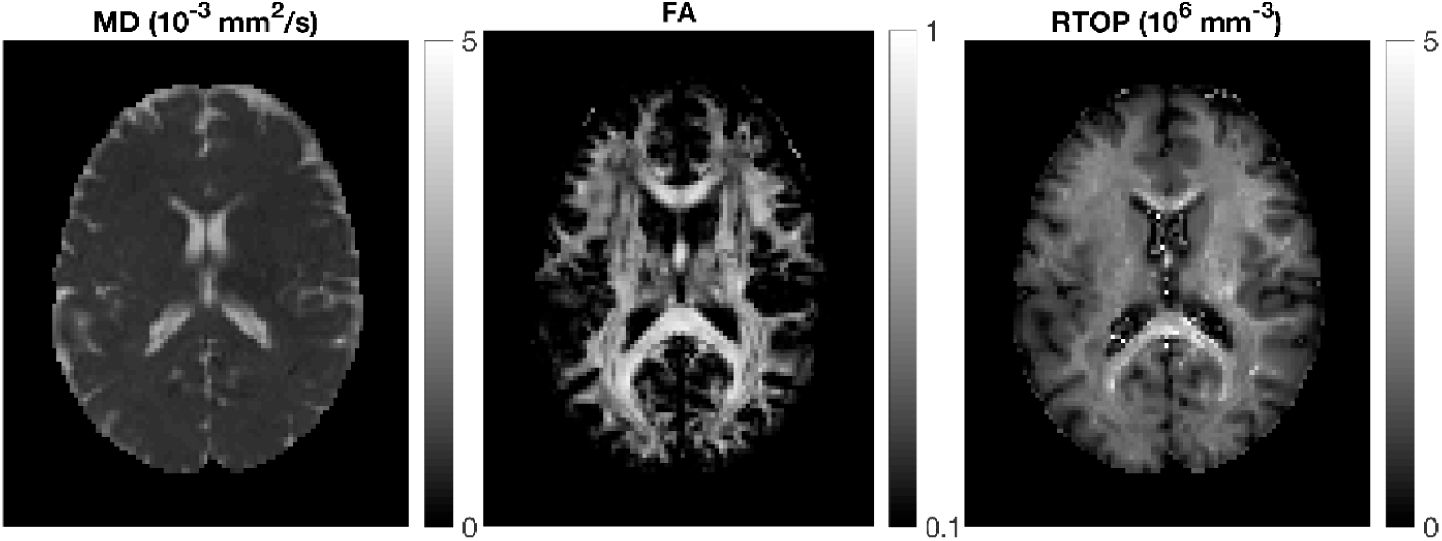
MD, FA and RTOP for subject MGH-1010, slice 45.

**Fig. 13.**
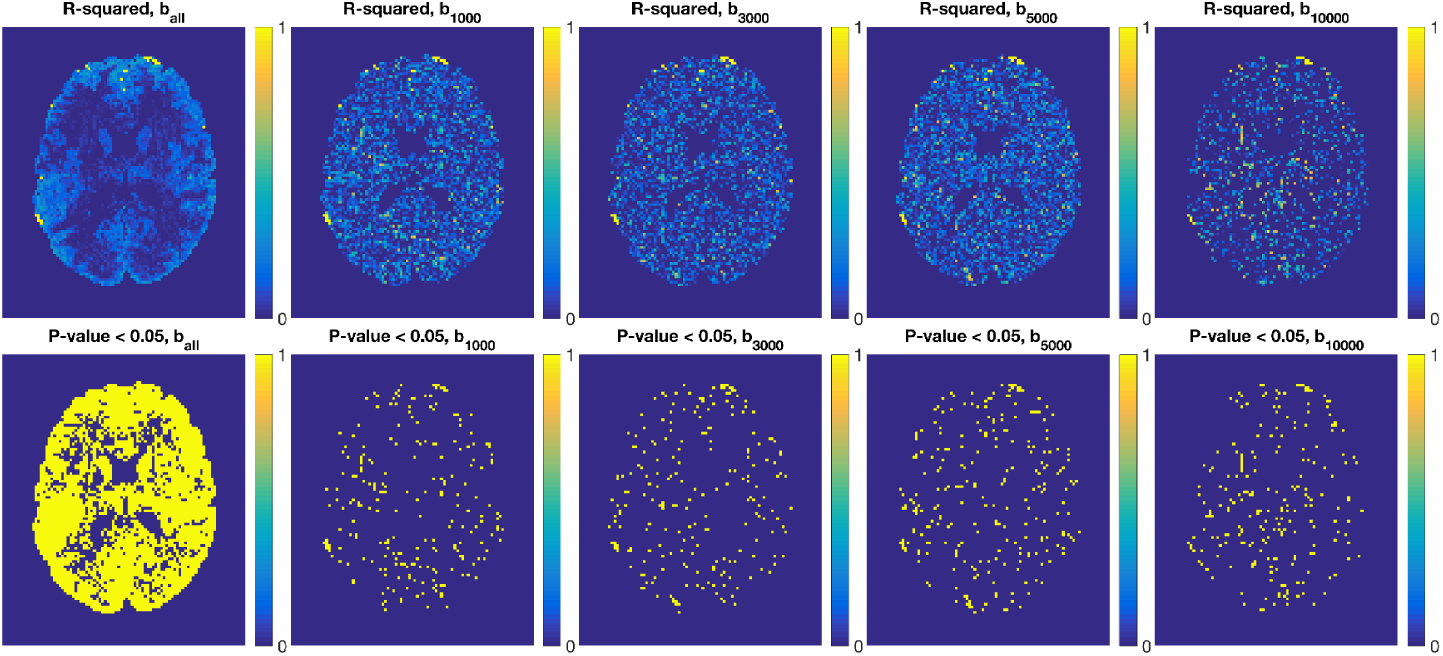
Results from the White test (testing for heteroscedastic variance); for all shells (column 1) and each shell separately (columns 2-5). R-squared values and voxels with a p-value ¡ 0.05 (uncorrected for multiple comparisons) are shown for subject MGH-1010, slice 45. The proportion of significant voxels in the brain mask are 5.1%, 4.9%, 5.3% and 5.0% for columns 2-5, i.e. close to the expected 5% for a threshold of p = 0.05.

#### Uncertainty of RTOP, RTAP and RTPP

In Figure 14, the standard deviation of RTOP for subjects MGH-1003, 1005, 1007 and 1010 are shown. There are two main clusters of voxels in the standard deviation maps wherein the white matter areas generally appear hyperintense, while the gray matter areas make up the lower intensity regions. There is limited contrast within the gray matter areas because it is relatively isotropic. Within the white matter areas, a large variability is detected in the corpus callosum areas for all four subjects. Within the corpus callosum, the splenium part shows a larger standard deviation than the genu and body of the corpus callosum.

**Fig. 14.**
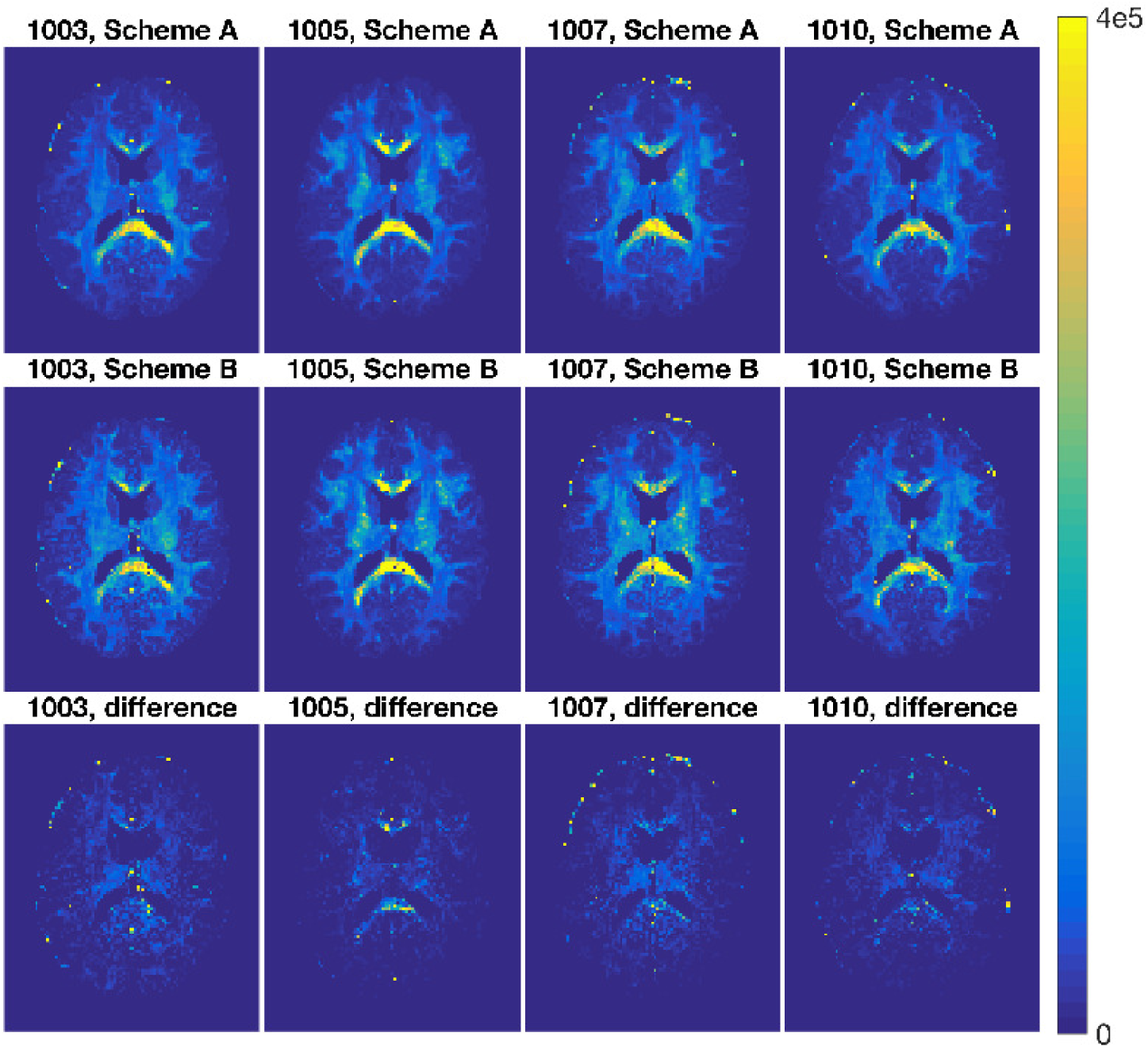
First and second rows: standard deviation of RTOP for subjects MGH-1003, 1005, 1007, 1010, slice 45, using two bootstrap sampling schemes and 500 bootstrap samples. Third row: absolute difference. Scheme A: sampling using all shells, Scheme B: sampling within each shell.

Figure 15 shows the QC of RTOP for subjects MGH-1003, 1005, 1007 and 1010. The QC measures dispersion of the distribution for 500 bootstrap RTOP samples without considering the RTOP values. Compared with the standard deviation maps in Figure 14, it is obvious that the contrast between the white matter and the gray matter decreases. A portion of the white matter regions shows larger QC values, most notably in the splenium of the corpus callosum. A larger dispersion is found in some gray matter voxels and this information and contrast are not available in the corresponding standard deviation maps. It is interesting to note the corpus callosum which has a highly anisotropic, coherent single fiber architecture demonstrates a much larger variability than other white matter areas. Figure 16 and 17 show the standard deviation and QC of RTAP and RTPP for subjects MGH-1003, 1005, 1007 and 1010, using bootstrap sampling Scheme A. Figure 18 shows the histograms of RTOP for a GM voxel and a WM voxel from subjects MGH-1003, 1005, 1007 and 1010.

**Fig. 15.**
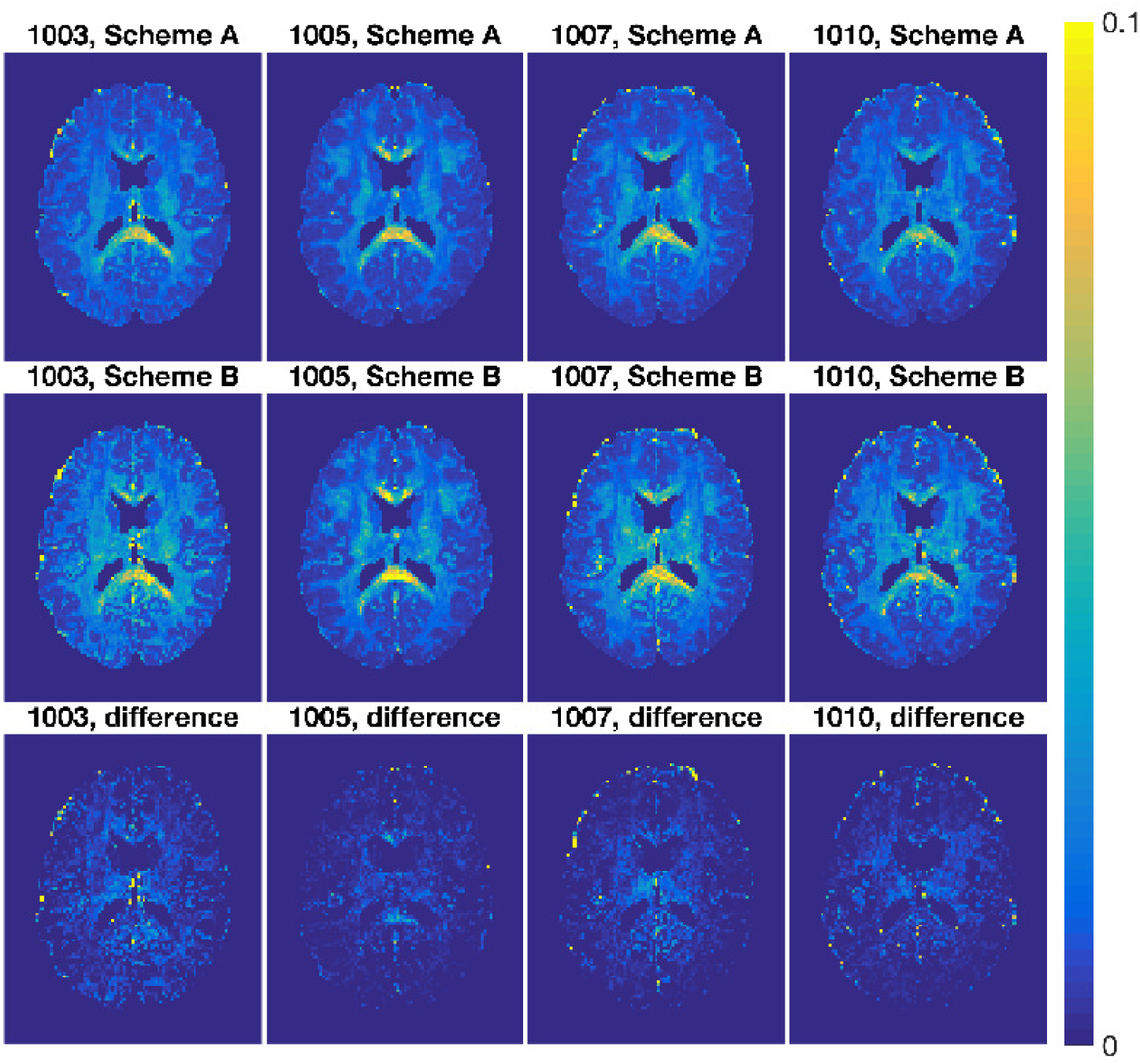
First and second rows: QC of RTOP for subjects MGH-1003, 1005, 1007, 1010, slice 45, using two bootstrap sampling schemes and 500 bootstrap samples. Third row: absolute difference. Scheme A: sampling using all shells, Scheme B: sampling within each shell.

**Fig. 16.**
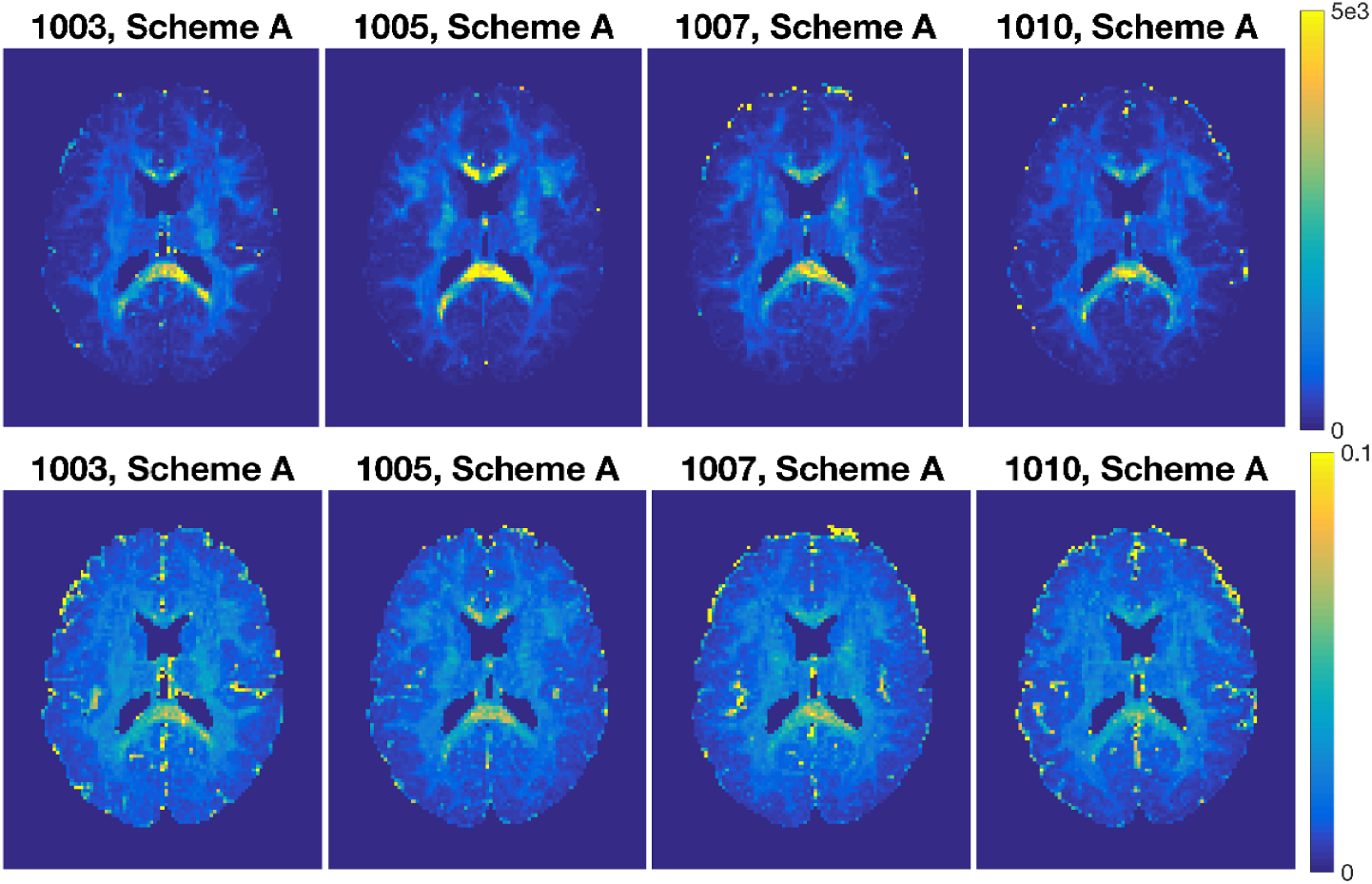
Std (top) and QC (below) of RTAP for subjects MGH-1003, 1005, 1007, 1010, slice 45, using bootstrap sampling Scheme A (sampling using all shells) and 500 bootstrap samples.

**Fig. 17.**
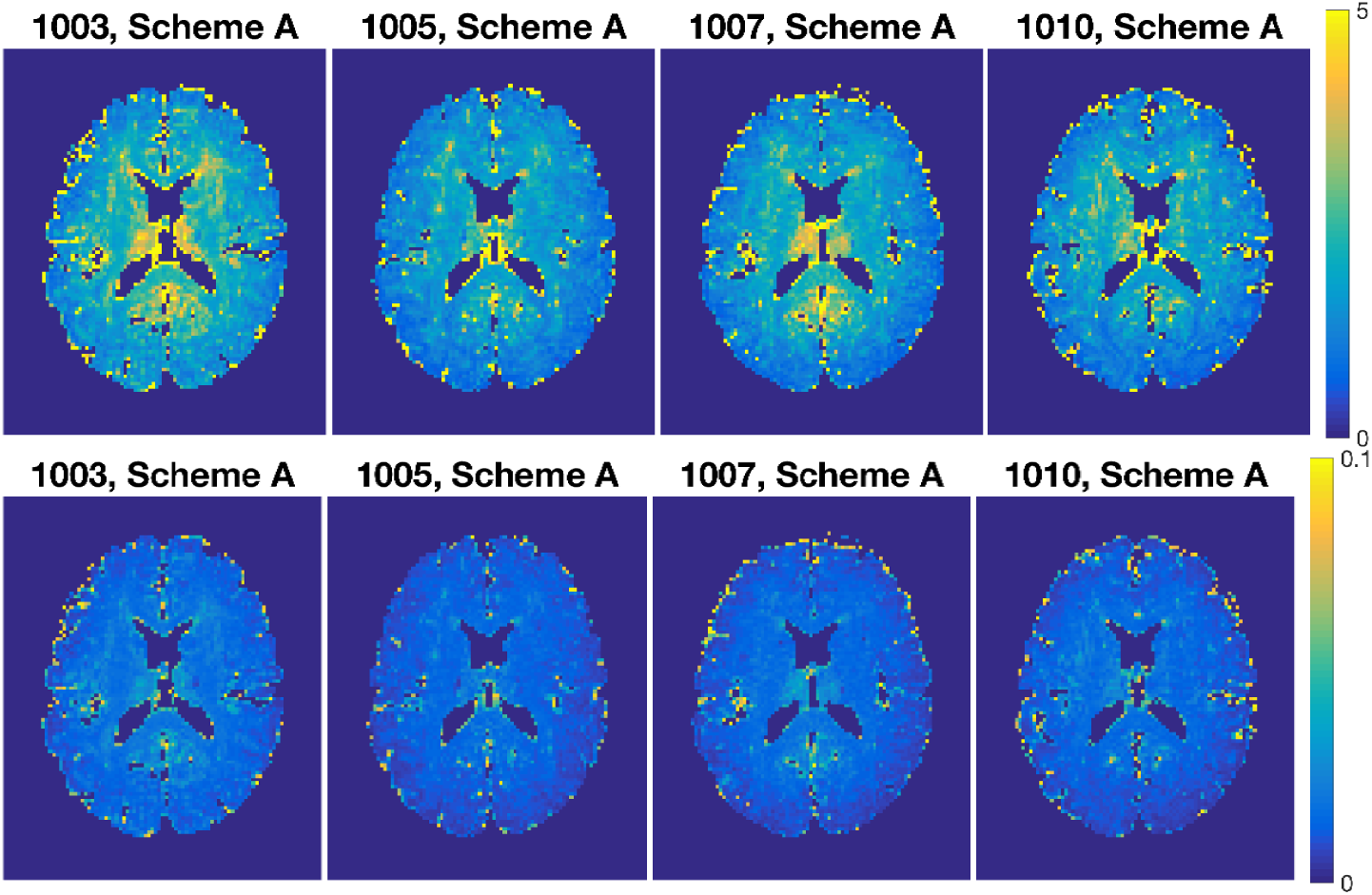
Std (top) and QC (below) of RTPP for subjects MGH-1003, 1005, 1007, 1010, slice 45, using bootstrap sampling Scheme A (sampling using all shells) and 500 bootstrap samples.

**Fig. 18.**
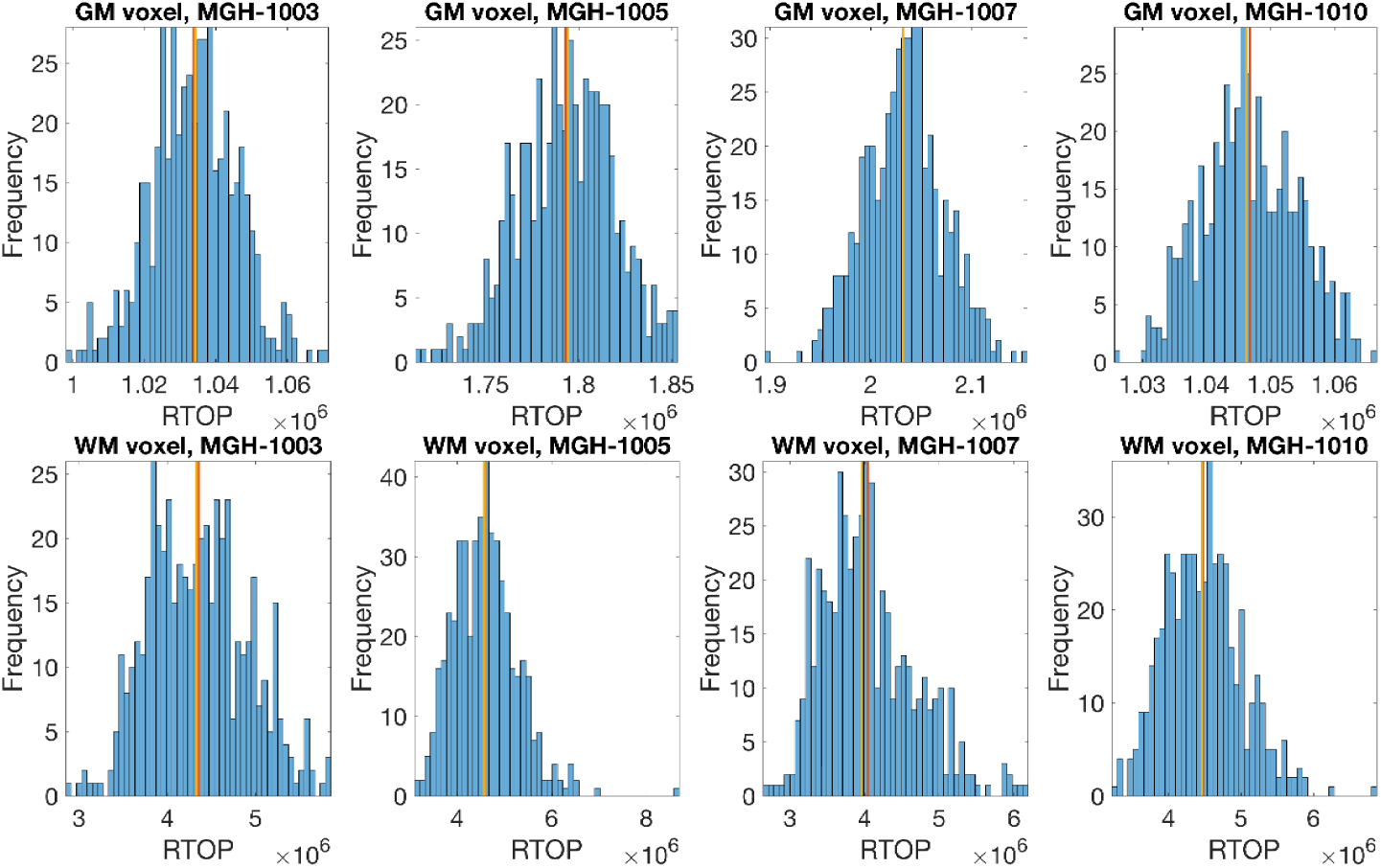
Histograms of RTOP for a GM voxel and a WM voxel, subjects MGH-1003, 1005, 1007, 1010, slice 45, using bootstrap Scheme A (sampling using all shells) and 500 bootstrap samples.

To study the uncertainty of diffusion metrics derived from MAP-MRI for different white matter tracts, we summarize the mean standard deviation and mean QC of RTOP for JHU ICBM-DTI-81 White-Matter Labels (Mori et al., 2005) in Appendix Table 1, using bootstrap Scheme A and 500 bootstrap samples. The JHU ICBM-DTI-81 White-Matter Labels are displayed in Figure 20. Both the standard deviation and the QC in Appendix Table 1 emphasize that the splenium of the corpus callosum exhibits relatively larger variability for the 500 bootstrap samples of RTOP, which is also accentuated in Figure 14 and 15. Splenium is the thickest part of the corpus callosum which connects the posterior cortices with fibers varying in size. For subjects 1003, 1005, and 1007, the splenium of the corpus callosum produces the largest QC values, and for subject 1010 the fornix gives the largest uncertainty. The mean QC of the anterior corona radiata tracts is substantially smaller than other labelled parts for all four subjects. However, in the regions where fibers cross or merge and the anisotropy becomes low, the uncertainty is seen to be high. Tractography results obtained for tracts that pass through such regions are likely to be very irreproducible and it would therefore be interesting to perform repeatability studies of tract reconstructions using MAP-MRI for tracts in these regions.

**Table 1.**
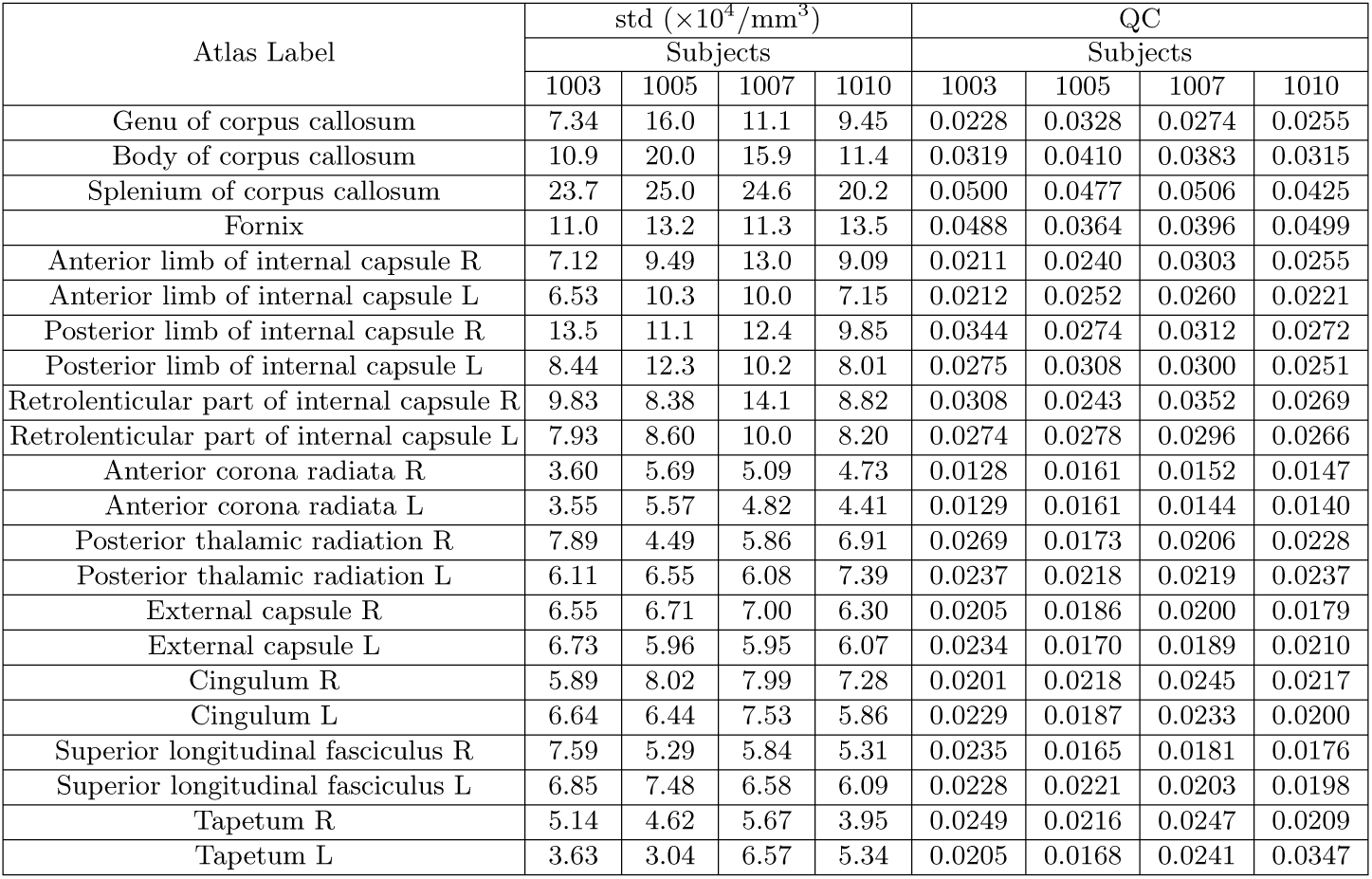
Mean std and QC of RTOP, JHU ICBM-DTI-81 White-Matter Labels, subject 1003, 1005, 1007 and 1010, slice 45, using bootstrap Scheme A (sampling using all shells) and 500 bootstrap samples.

Appendix Table 2 shows the mean standard deviation and mean QC of RTOP for JHU ICBM-DTI-81 White-Matter Labels using bootstrap Scheme B and 500 bootstrap samples. It is notable that sampling residuals within each shell leads to slightly larger variability for all JHU white matter tracts. Bootstrap sampling serves primarily as a measure of precision for diffusion-derived metrics, so there is the expected correlation with the SNR value. It is reported in (Vorburger et al., 2016) that the dispersion of diffusion tensor principal eigenvector in fornix is correlated with the subjects’ memory performance. As expected, the splenium of the corpus callosum produces relatively larger standard deviation and QC values, and for subjects 1003, 1007 and 1010, the fornix tract demonstrates the largest variability. The anterior corona radiata tracts always produce the least uncertainty, regardless of the bootstrap sampling schemes using all shells or within the same shell.

**Table 2.**
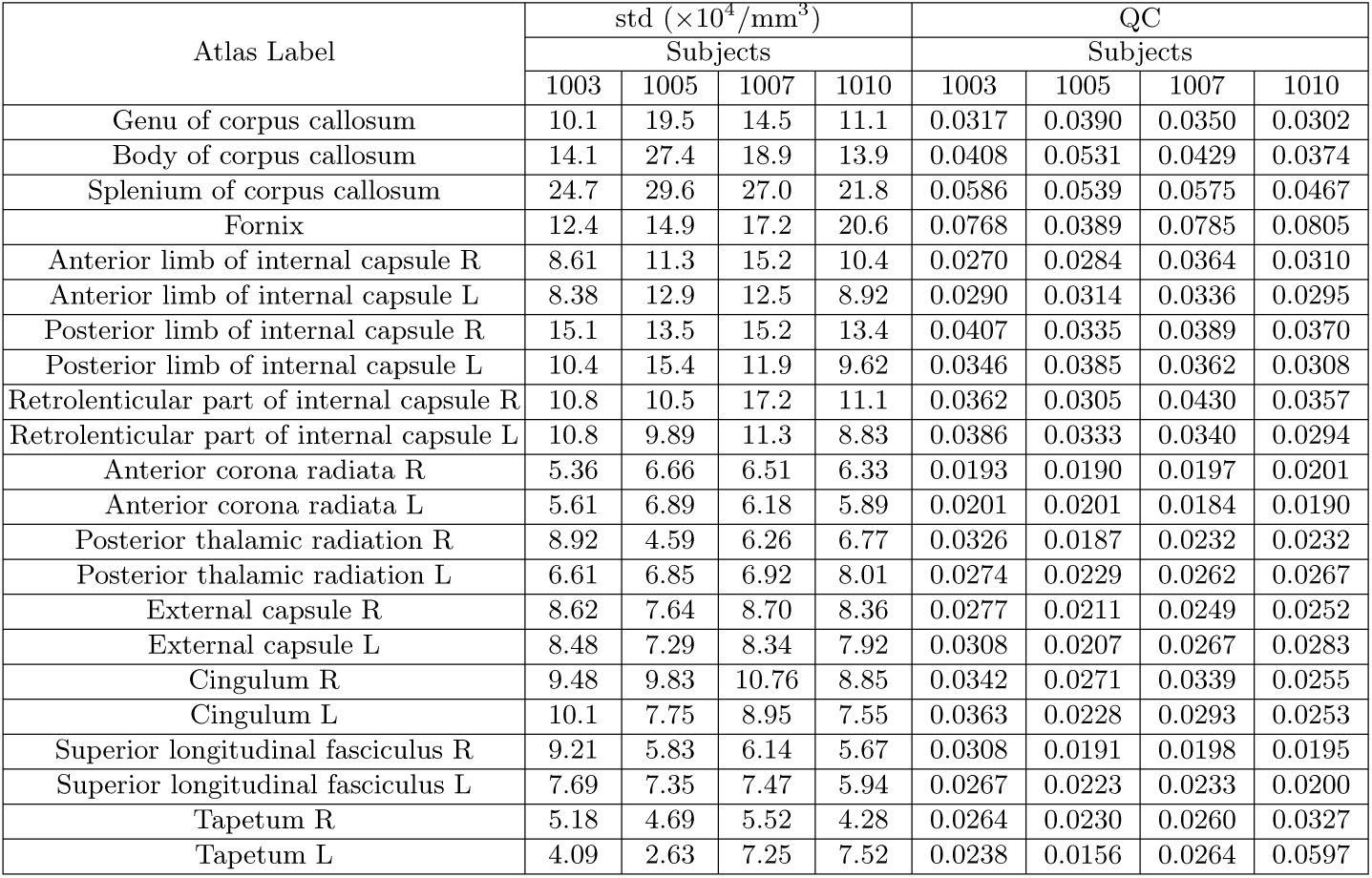
Mean std and QC of RTOP, JHU ICBM-DTI-81 White-Matter Labels, subject 1003, 1005, 1007 and 1010, slice 45, using bootstrap Scheme B (sampling within the shell) and 500 bootstrap samples.

Appendix Table 3 shows the mean QC of RTAP and RTPP for JHU ICBM-DTI-81 White-Matter Labels using bootstrap Scheme A and 500 bootstrap samples. The best precision is achieved by the anterior corona radiata, followed by superior long fasciculus for RTAP. Generally, the statistics of RTAP demonstrates a similar behavior as RTOP. The RTPP and RTAP values can be seen as the “decomposition” of the RTOP values into components parallel and perpendicular to the direction of the primary eigenvector of the diffusion tensor, respectively. Generally, the variation pattern across the white matter tracts is similar for all three investigated MAP-MRI metrics. In other words, if the uncertainty in a white matter tract is high with respect to RTOP, it is also high with respect to the other two parameters, i.e., RTAP and RTPP. Appendix Tables 1, 2 and 3 indicate that we should be less confident in the MAP-MRI derived metrics from those tracts with relatively higher QC values.

**Table 3.** Mean QC of RTAP and RTPP, JHU ICBM-DTI-81 White-Matter Labels, subject 1003, 1005, 1007 and 1010, slice 45, using bootstrap Scheme A (sampling using all shells) and 500 bootstrap samples.

| Atlas Label | RTAP |  |  |  | RTPP |  |  |  |
| --- | --- | --- | --- | --- | --- | --- | --- | --- |
|  | Subjects |  |  |  | Subjects |  |  |  |
|  | 1003 | 1005 | 1007 | 1010 | 1003 | 1005 | 1007 | 1010 |
| Genu of corpus callosum | 0.0260 | 0.0329 | 0.0276 | 0.0275 | 0.0211 | 0.0158 | 0.0166 | 0.0173 |
| Body of corpus callosum | 0.0324 | 0.0388 | 0.0357 | 0.0305 | 0.0317 | 0.0235 | 0.0249 | 0.0253 |
| Splenium of corpus callosum | 0.0482 | 0.0432 | 0.0467 | 0.0396 | 0.0226 | 0.0190 | 0.0193 | 0.0182 |
| Fornix | 0.0567 | 0.0362 | 0.0617 | 0.0358 | 0.0461 | 0.0317 | 0.0425 | 0.0354 |
| Anterior limb of internal capsule R | 0.0267 | 0.0247 | 0.0308 | 0.0253 | 0.0181 | 0.0143 | 0.0154 | 0.0156 |
| Anterior limb of internal capsule L | 0.0256 | 0.0250 | 0.0276 | 0.0249 | 0.0184 | 0.0147 | 0.0165 | 0.0186 |
| Posterior limb of internal capsule R | 0.0330 | 0.0262 | 0.0298 | 0.0258 | 0.0168 | 0.0136 | 0.0152 | 0.0151 |
| Posterior limb of internal capsule L | 0.0285 | 0.0291 | 0.0287 | 0.0252 | 0.0179 | 0.0147 | 0.0167 | 0.0169 |
| Retrolenticular part of internal capsule R | 0.0299 | 0.0250 | 0.0347 | 0.0268 | 0.0198 | 0.0155 | 0.0174 | 0.0164 |
| Retrolenticular part of internal capsule L | 0.0285 | 0.0265 | 0.0284 | 0.0283 | 0.0186 | 0.0151 | 0.0157 | 0.0164 |
| Anterior corona radiata R | 0.0207 | 0.0199 | 0.0187 | 0.0191 | 0.0220 | 0.0151 | 0.0158 | 0.0166 |
| Anterior corona radiata L | 0.0197 | 0.0189 | 0.0195 | 0.0203 | 0.0210 | 0.0153 | 0.0176 | 0.0182 |
| Posterior thalamic radiation R | 0.0279 | 0.0178 | 0.0220 | 0.0232 | 0.0189 | 0.0144 | 0.0160 | 0.0141 |
| Posterior thalamic radiation L | 0.0262 | 0.0225 | 0.0247 | 0.0245 | 0.0220 | 0.0141 | 0.0172 | 0.0156 |
| External capsule R | 0.0244 | 0.0201 | 0.0214 | 0.0224 | 0.0187 | 0.0142 | 0.0151 | 0.0164 |
| External capsule L | 0.0263 | 0.0194 | 0.0218 | 0.0239 | 0.0181 | 0.0164 | 0.0173 | 0.0191 |
| Cingulum R | 0.0232 | 0.0248 | 0.0263 | 0.0234 | 0.0238 | 0.0162 | 0.0180 | 0.0170 |
| Cingulum L | 0.0266 | 0.0228 | 0.0250 | 0.0212 | 0.0200 | 0.0152 | 0.0180 | 0.0168 |
| Superior longitudinal fasciculus R | 0.0256 | 0.0197 | 0.0212 | 0.0201 | 0.0164 | 0.0146 | 0.0153 | 0.0146 |
| Superior longitudinal fasciculus L | 0.0248 | 0.0216 | 0.0214 | 0.0205 | 0.0171 | 0.0119 | 0.0142 | 0.0134 |
| Tapetum R | 0.0291 | 0.0232 | 0.0253 | 0.0333 | 0.0382 | 0.0347 | 0.0332 | 0.0262 |
| Tapetum L | 0.0250 | 0.0193 | 0.0250 | 0.0430 | 0.0385 | 0.0376 | 0.0298 | 0.0261 |

#### Group analysis

Similar to functional MRI it is common to perform group studies using diffusion MRI, to for example find differences between healthy controls and subjects with some disease. One of the most common scalar measures for group analysis is fractional anisotropy, calculated from the diffusion tensor, which for example has been used to study diffuse axonal injuries in mild traumatic brain injury (Eierud et al., 2014; Shenton et al., 2012). A weighted mean can be calculated as

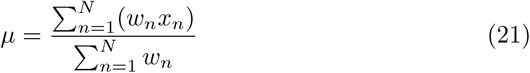

where 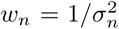. Instead of each voxel value contributing equally to the final mean, voxels with higher standard deviation contribute less “weight” than others. A comparison between the mean RTOP and the weighted mean RTOP is presented in Figure 19. The weighted mean for example downweights an outlier close to the posterior cingulate. In Figure 19, it is also presented a comparison of the unweighted and the weighted probability density distributions of RTOP^1/3^ for one voxel. Subjects with a higher uncertainty will be downweighted, which can lead to a different distribution of the mean RTOP^1/3^.

**Fig. 19.**
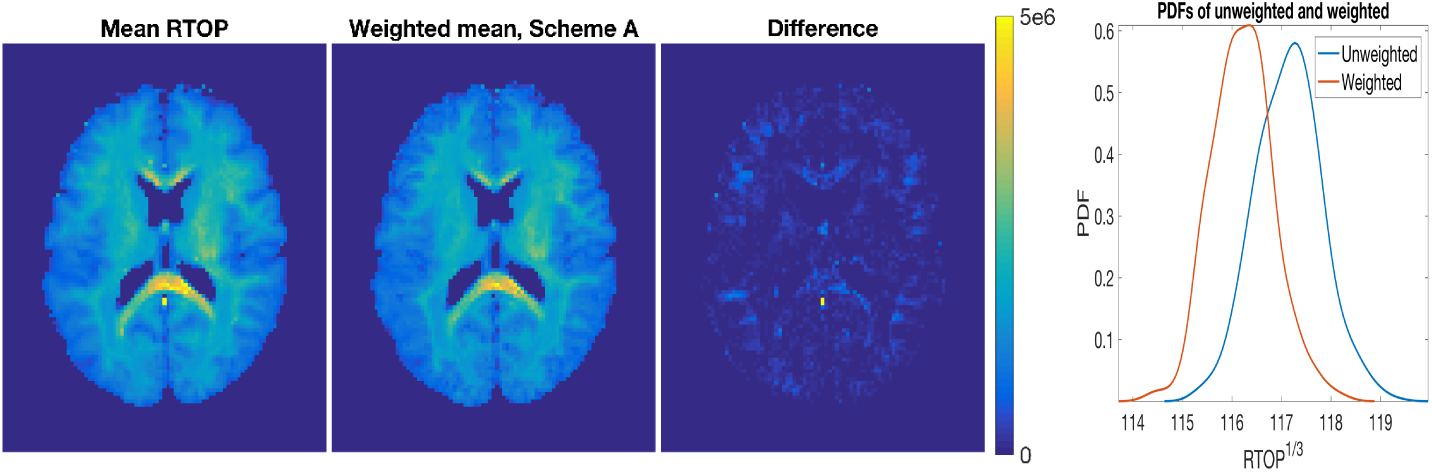
Mean and weighted mean of RTOP, PDFs of unweighted and weighted RTOP, subjects MGH-1003, 1005, 1007, 1010, slice 45, using bootstrap Scheme A (sampling using all shells) and 500 bootstrap samples.

**Fig. 20.**
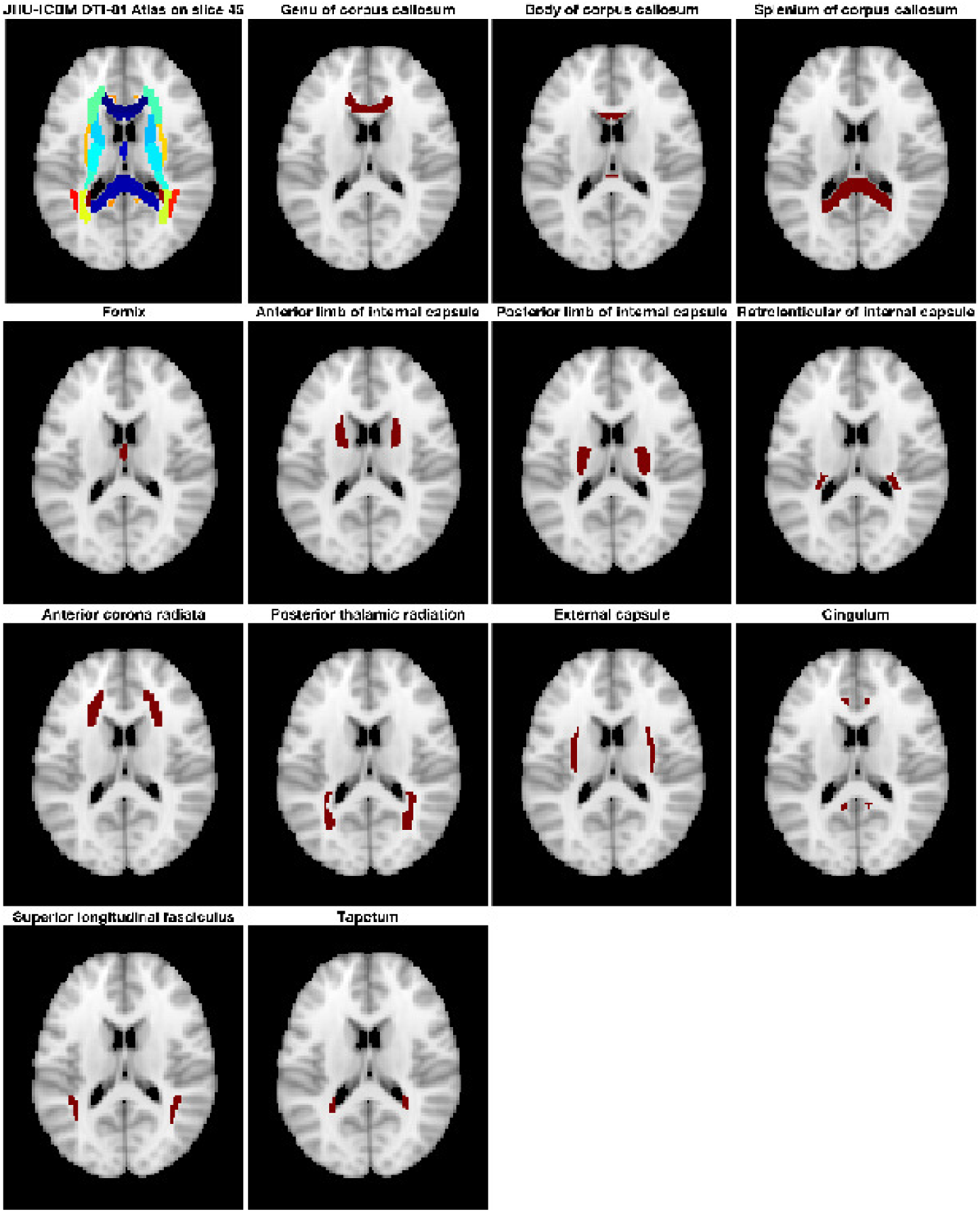
JHU-ICBM DTI-81 Atlas on slice 45.

## 5 Discussion

Researchers have previously applied the bootstrap method to the residuals from the linear regression used to fit the diffusion tensor model. We have extended this technique to a more sophisticated diffusion model to quantify the uncertainty of its derived metric. We used the residual bootstrap technique to evaluate the uncertainty of the MAP-MRI derived metrics using physical phantom data as well as human brain data. To the best of our knowledge, this is the first work that investigates the uncertainty of diffusion metrics derived from MAP-MRI.

The experiments are divided into two sections: one dealing with uncertainty estimation for physical phantom data in which four sets of data are collected using different number of measurements and different b-values, and the other dealing with uncertainty estimation for human brain data of four healthy subjects collected using the same scan protocol. The variability originates from measurement noise and is influenced by a wide range of parameters, many of which are difficult or impossible to fully model. It is generally assumed that diffusion data are primarily affected by thermal, normally distributed noise. However, physiological noise and artifacts may also affect diffusion data and may result in non-normal and spatially variant noise characteristics.

Several studies report that for the diffusion tensor model, the variability in the tensor trace, diffusion anisotropy, and the tensor major eigenvector are related to the spatial orientation of the tensor (Batchelor et al., 2003; Jones and Pierpaoli, 2004). The orientational dependence of the tensor variance decreases for both more uniformly distributed encoding schemes and increased number of encoding directions. In the experiment using SPARC data, we have demonstrated that the variability of MAP-MRI derived metrics decrease for both increased number of shells and increased number of gradient directions in each shell. The variation in single-fiber area is less sensitive to the number of gradient directions on each shell. A voxel with two fiber bundles and a voxel with a single fiber bundle might lead to the same RTOP value, but the uncertainty might be affected differently. It can be concluded that using a large number of shells is more efficient to minimize the variation in the MAP-MRI diffusion metrics, compared with a large number of measurements on the same shell.

Bootstrap Schemes A and B, sampling residuals using all shells and within each shell, produce essentially identical results of standard deviation and QC when applied to the SPARC data. This may be because that in the SPARC data there is only one measurement for the shell *b*_0_. It should be noted that when using bootstrap Scheme B, that is sampling residuals within each shell, an increased variability is found for MAP-MRI metrics in the HCP-MGH human brain diffusion data (Figure 14 and 15) but is not found in the SPARC data.

The probability densities of RTOP were estimated to investigate configuration of local diffusion profiles in the SPARC data. It is demonstrated that the probability density follows expected patterns corresponding to the fiber configurations in the physical phantom using both bootstrap sampling schemes. These patterns do not differ much when the number of gradient directions on each shell is changed, but provide more details when more shells are used in the diffusion scan protocols.

It is important to note that all acquisition parameters which influence the SNR of the diffusion signals, such as the b-value, the number of measurements, the gradient strength, the echo time, etc., most likely have a direct influence on the uncertainty.

In (Avram et al., 2016), the computation time for the reconstruction of MAP-MRI parameters from whole-brain diffusion data sets (70 × 70 × 42 × 698) using *N*_max_ = 6 was less than 3 h on a single workstation with 32GB RAM and 8 cores (Intel i7-4770 K at 3.5G Hz). In (Fick et al., 2016), it is reported that it takes around 60 seconds to do the MAP-MRI fitting (*N*_max_ of 6) for all voxels of SPARC-30, using an Intel(R) Core(TM) i7-3840QM CPU with 32 GB RAM. In this paper, we use two Intel(R) Xeon(R) E5-2697 CPUs and OpenMP to support multi-thread processing, which makes it possible to do the MAP-MRI fitting (*N*_max_ of 6) for all voxels of SPARC-30 within 2 seconds. To run 500 bootstrap replicates takes about 40 minutes and 20 hours, respectively for the SPARC data and a slice of the HCP-MGH data. In theory, graphics processing units (GPUs) can be used for further speedup (Eklund et al., 2013, 2014) as they can process some 30 000 voxels in parallel.

In conclusion, bootstrap metrics, such as the standard deviation and QC, provide additional valuable diffusion information next to the common MAP-MRI parameters and should be incorporated in ongoing and future MAP-MRI studies to provide more extensive insight.

## Acknowledgements

This research is supported by the Swedish Research Council (grant 2015-05356, “Learning of sets of diffusion MRI sequences for optimal imaging of micro structures”), Link¨oping University Center for Industrial Information Technology (CENIIT), and the Knut and Alice Wallenberg Foundation project “Seeing Organ Function”.

Data collection and sharing for this project is provided by the Human Connectome Project (HCP; Principal Investigators: Bruce Rosen, M.D., Ph.D., Arthur W. Toga, Ph.D., Van J. Weeden, MD). HCP funding was provided by the National Institute of Dental and Craniofacial Research (NIDCR), the National Institute of Mental Health (NIMH), and the National Institute of Neuro-logical Disorders and Stroke (NINDS). HCP data are disseminated by the Laboratory of Neuro Imaging at the University of Southern California.

## A Appendix

